# Systematic prediction and functional analysis of amino acid residues determining product specificity in the plant oxidosqualene cyclase superfamily

**DOI:** 10.64898/2026.05.26.727834

**Authors:** Rashmi Kumari, Neeladri Sen, Rebecca Casson, Charlotte Owen, Michael J Stephenson, Neera Borkakoti, Christine Orengo, Janet M. Thornton, Anne Osbourn

**Affiliations:** European Bioinformatics Institute (EMBL-EBI), Wellcome Genome Campus, Hinxton, CB10 1SD, Cambridgeshire, UK; Institute of Structural and Molecular Biology, University College London, London WC1E 6BT, UK; Department of Biochemistry and Metabolism, John Innes Centre, Norwich Research Park, Norwich NR4 7UH, UK; School of Chemistry, Pharmacy and Pharmacology, University of East Anglia, Norwich Research Park, Norwich NR4 7TJ, UK

**Author notes:** Correspondence to: Christine Orengo,; Janet Thornton,; Anne Osbourn. DIOSynVax Ltd, Department of Veterinary Medicine, University of Cambridge, Madingley Road, Cambridge, CB3 0ES, UK. These authors contributed equally to this work (Rashmi Kumari, Neeladri Sen).

## Abstract

Oxidosqualene cyclases (OSCs) catalyse one of nature’s most intricate enzyme reactions, converting the linear precursor 2,3-oxidosqualene into an array of cyclic triterpene scaffolds through sequential carbocation cascades. Predicting OSC function based on sequence is challenging beyond broad family-level classification. Here, we develop a structure-based computational framework to identify amino acid determinants of OSC product specificity. Using 169 functionally characterised OSCs, we deploy a multifaceted approach combining differential conservation along with structural information, physico-chemical properties of amino acids and binding pocket electrostatics in order to understand the determinants of product specificity. Using *Arabidopsis thaliana* cycloartenol synthase AtCAS as a model, we then validate our predictions through targeted mutagenesis, achieving stepwise reprogramming towards the protosteryl-type products cucurbitadienol and lanosterol, including complete product switches. Molecular dynamics simulations support a mechanism in which subtle pocket remodelling alters active-site volume, water access and proton-elimination chemistry. These findings provide a blueprint for OSC engineering.

## Introduction

Computational methods for predicting protein function from sequence are well explored. The regular Comparative Assessment of Function Annotation (CAFA) exercise^1,2^ assesses the accuracy of function prediction methods that aim to annotate a novel sequence with its molecular function, biological process and cellular location. Sequences are distributed to participants, who provide a computational prediction of the function (expressed as Gene Ontology terms or Human Phenotype Ontology terms), and the results are compared to experimental data collected worldwide for individual proteins. Many of the prediction methods available include components based on finding evolutionary relationships with proteins of known function^3–5^. These approaches are excellent for broad classifications, for example in distinguishing an oxidoreductase and a transferase^1,2^, but are less accurate in differentiating the specific functions of individual members in an enzyme family. Most enzyme families evolve to perform different functions by modulating the substrate on which they operate. Prediction therefore usually involves docking different substrates into the enzyme active sites to assess how well they bind. The situation is entirely different, however, for enzyme families such as oxidosqualene cyclases (OSCs; also known as triterpene synthases), which typically use the same starting substrate (2,3-oxidosqualene) but generate a diverse array of products^6–10^.

The terpenes are the largest and most structurally diverse family of natural products, with over 80,000 reported to date^11^. Their scaffolds are biosynthesised from linear substrates containing multiple 5-carbon (C-5) isoprene units to give cyclized molecules of varying complexity, e.g. monoterpenes (C-10), sesquiterpenes (C-15), diterpenes (C-20) and triterpenes (C-30), of which the triterpenes are the most complex^12^. Triterpene-derived metabolites are particularly highly diversified in plants (>20,000 reported to date)^12^ and have important functions in nature, providing protection against pests and pathogens, shaping the root microbiome, and influencing crop quality^13–15^. They are also precursors for essential steroid membrane components and hormones, and are of considerable interest for their pharmaceutical properties^16^.

The initial cyclisation and subsequent cationic rearrangement of 2,3-oxidosqualene by OSCs is arguably the most complex single enzyme transformation observed in nature^17^. A single catalytic residue (an aspartic acid; ASP) triggers catalysis by protonation of the terminal epoxide of 2,3-oxidosqualene (Fig. 1). The resulting carbocation then undergoes a series of electrophilic additions to proximal alkenes, eventually yielding a tetracyclic cationic intermediate. Resolution of this intermediate typically involves further carbocation migration via cascades of 1,2-shifts, with or without additional ring annulations. The reaction is usually terminated either via the elimination of a neighboring hydrogen to give a neutral alkene or by the quenching of the terminal cation by water to give a diol. More complex rearrangements may also occur leading to a plethora of different products from a single substrate (e.g. over 200 distinct cyclisation products for 2,3-oxidosqualene^18^), mandated by the environment of the substrate in the enzyme binding pocket.

**Fig. 1.**
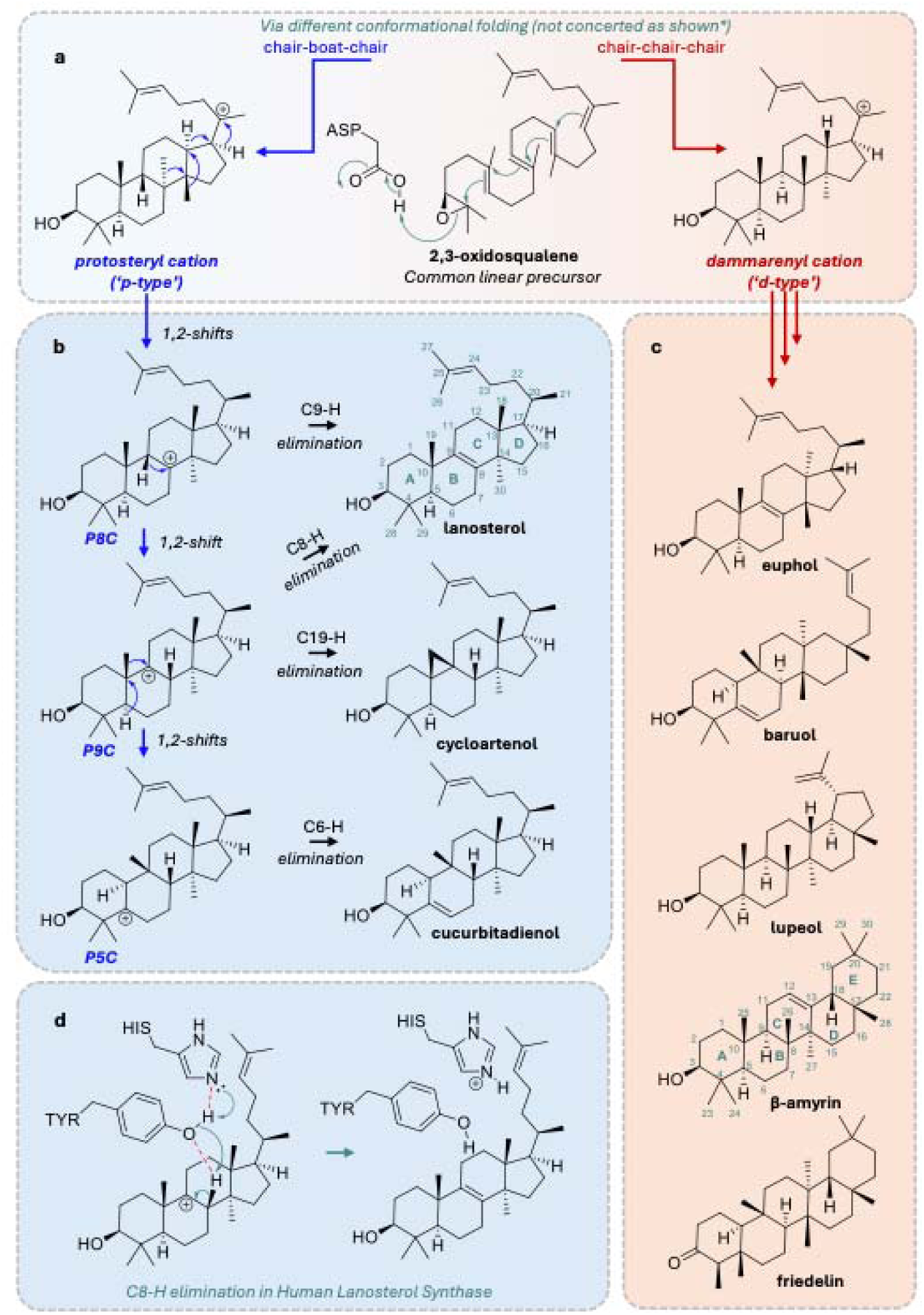
Biosynthetic origin of triterpene scaffolds from the common precursor 2,3-oxidosqualene. **a**, Established dichotomy between a protosteryl and dammarenyl cation origin; * initial cyclisation is likely to progress through intermediates of differing ring numbers. **b**, Reaction pathway relationships between the protosteryl-derived triterpenes lanosterol, cycloartenol, and cucurbitadienol. Tetracyclic triterpene carbon numbering and ring nomenclature are given for lanosterol. Key C9, C8, and C5 carbocations are highlighted and labelled P9C, P8C, and P5C, respectively. **c,** Examples of dammarenyl-derived triterpenes. Pentacyclic triterpene carbon numbering and ring nomenclature are given for β-amyrin. **d,** Tyrosine-histidine catalytic dyad responsible for C8-H elimination to give lanosterol in human lanosterol synthase^47^.

Whilst numerous mutational studies have been carried out on OSCs, these have invariably been done on a case-by-case basis to study a given enzyme or pathway of interest^10,18–22^. Prediction of the function of OSCs based on sequence thus represents a challenging computational problem. Here we use computational approaches to deploy a systematic analysis of the complex relationship between structure and function for this enzyme superfamily. In particular, to detect function-determining residues we employ a strategy that identifies residue positions that are differentially conserved between diverse protein subfamilies (functional families - in which all relatives have the same functional properties). This approach (FunFamer^23^) exploits the statistically robust Groupsim method^24^, has successfully illuminated key residues determining specificity and functional mechanism in other enzyme families^25^, and been endorsed by the CAFA evaluation^26^. We further identify and experimentally validate product specificity-determining residues that determine the outcome of cyclization processes for OSCs that generate protosteryl-type products (Fig. 1b) and shed light on the underlying mechanisms of triterpene cyclization.

## Results

### *In silico* analysis of characterized OSC enzymes

To predict and map genome-encoded triterpene scaffold diversity across the plant kingdom we first collated a set of 169 characterized plant OSC sequences, including committed OSCs that make a single product and others that make more than one product (multifunctional OSCs) (Data S1). Sequence similarity network (SSN) and phylogenetic analysis of these functionally validated OSCs were consistent with the literature^6,10,19–22^ (Fig. 2a). There was a clear distinction between the two major groups of enzymes that make protosteryl-type (‘p-type’) products and dammarenyl-type (‘d-type’) products, independent evolution of d-type products, and some conservation in sequence for specific products such as lupeol and friedelin (Fig. 2a)^9,10^. Within the p-type OSCs, the three primary groups are cycloartenol, lanosterol and cucurbitadienol synthases. For the d-type enzymes, there is a less well-defined relationship between sequence and function. The most frequently characterized function is β-amyrin synthesis, which is carried out by committed and multi-functional OSCs across the Plant Kingdom. However, these OSCs do not form a monophyletic clade and instead group with other d-type products such as lupeol, baurenol and taraxerol (Fig. 2a)^10^. Overall, it can be observed that some d-type OSC clades appear to have relatively conserved functions, such as the friedelin synthases, whereas other functions have independently evolved multiple times, such as β-amyrin and lupeol synthesis^19,20^. A clear understanding of the reaction mechanisms of OSCs is therefore necessary in order to unpick the intricacies of the sequence-to-function relationship of this enzyme family.

**Fig. 2.**
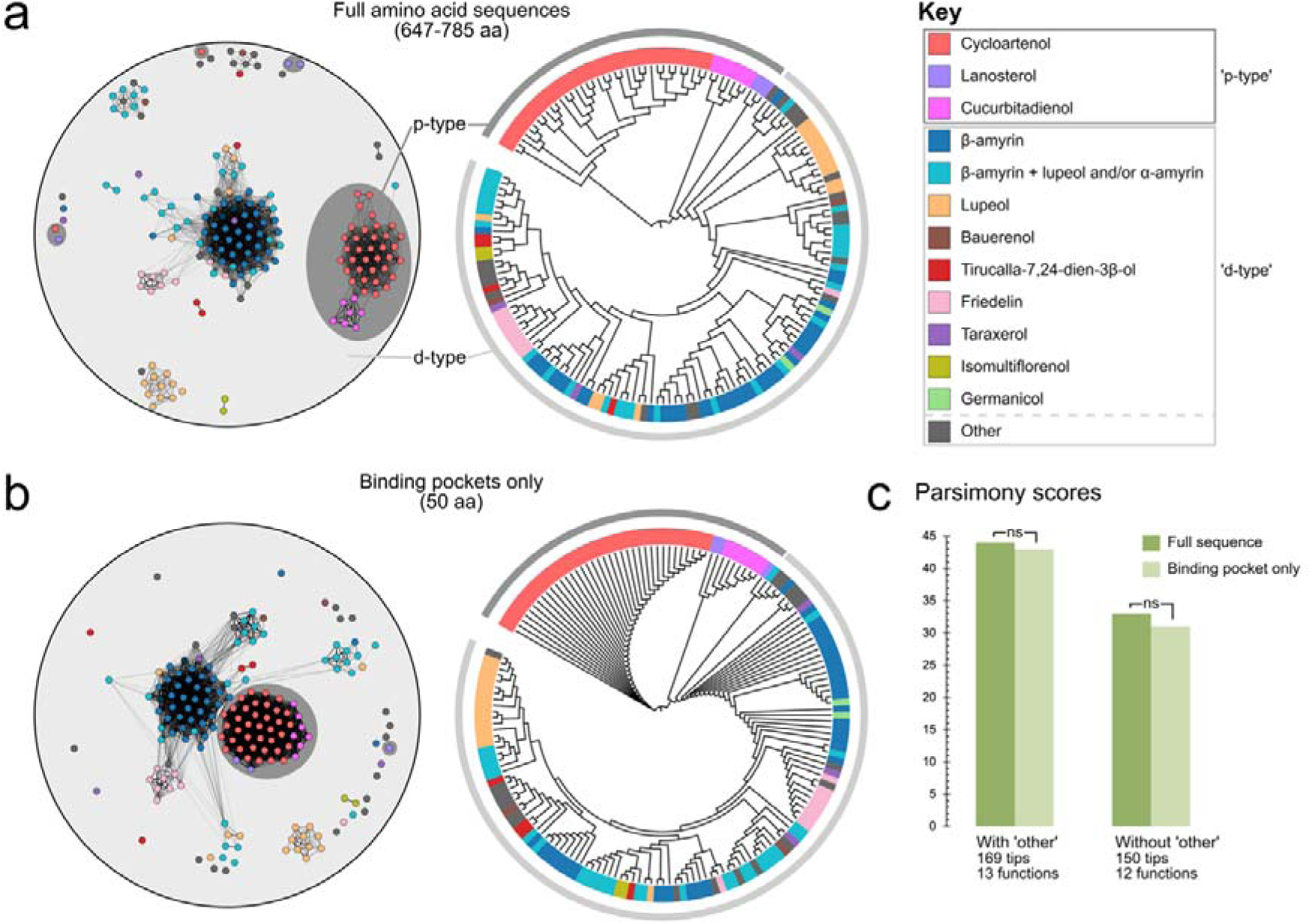
Clustering and phylogeny of functionally characterized OSCs. a, Sequence similarity network (SSN; left) and cladogram (right) for the full amino acid sequences of the 169 characterised OSCs (Data S1). Products of the analysed OSCs are indicated by the colour according to the key, with ‘p-type’ products additionally highlighted in dark grey and ‘d-type’ products additionally highlighted by light grey. Multifunctional OSCs are labelled according to their primary product. ‘Other’ represents OSCs for which only one example of their primary product exists in the reference set. **b,** SSN and cladogram for the 169 OSCs using only the 50 amino acid residues constituting the predicted binding pocket (Methods, Data S2). Colours represent OSC products as in **a,** according to the key. **c,** Parsimony scores (assessed by Bayesian tip-association significance testing; Materials and Methods) for the cladograms shown in **a** and **b**. A lower score represents a closer link between sequence and function. When including or excluding all ‘other’ OSCs as a discrete functional group, no significant difference was calculated between the full length and binding pocket cladograms. For the full set of OSC sequences (169 tips and 13 functions) full length vs. binding pocket ΔPS =-2, one-sided *p* = 0.339, z =-0.591. Without sequences classified as ‘other’ (150 tips and 12 functions) full length vs. binding pocket ΔPS =-1, one-sided *p* = 0.451, z =-0.256.

The reaction mechanism can be divided into two major steps: firstly binding and folding of 2,3-oxidosqualene into a ring-like configuration, and secondly ring closures and carbocation rearrangement. A quantum molecular dynamics study for smaller terpenes (mono-and sesquiterpenes) has shown that ring formation (10^-9^ s) is significantly slower than carbocation rearrangement (10^-15^ s); also the timescale of the ring formation does not significantly change in the presence or absence of the terpene synthase enzyme^27^. By analogy, 2,3-oxidosqualene is expected to first adopt a ring-like configuration, as observed for squalene in the crystal structure of the bacterial squalene-hopene cyclase (SHC) enzyme^28^. Once the reaction begins, ring closures and carbocation rearrangement are likely to happen so quickly that only small local conformational changes will occur, i.e. the carbocation rearrangements may be so fast that they can only be influenced by the local conformation of the substrate, the neighbouring binding residues, or water molecules. Binding pocket residues are therefore likely to play a critical role in determining product specificity. We therefore investigated whether we could use binding pocket residues to better classify enzymes that generate different products into separate branches of committed and multifunctional groups.

Although a tetrameric cryo-EM structure has been generated for an OSC from the Chinese medicinal plant *Tripterygium wilfordii*^29^, there is as yet no high resolution, monomeric crystal structure available for a plant OSC with bound substrates/inhibitors. However, crystal structures are available for two other triterpene cyclases, namely, the OSC human lanosterol synthase (HLS, PDB ID: 1W6K)^30^, and the bacterial enzyme squalene hopene cyclase (SHC, PDB ID: 1UMP)^31,32^, which share around 25% amino acid sequence identity (Supplementary Table 1). A structure for SHC bound to the sterol biosynthesis inhibitor 2-azasqualene, an analogue of squalene, has also been determined^28^. We superimposed the structures of HLS and SHC and considered all amino acid residues within 5Å of 2-azasqualene in both enzymes. Additionally, if any residues of a loop were within 5Å, the whole loop was considered as binding pocket residues since the loops may be flexible and the residue positions could be slightly different for other enzymes. This resulted in a total of 50 amino acid residues (Supplementary Fig. 1). The predicted structures of the 169 characterised plant OSC sequences (Data S1) were downloaded from the AlphaFold Protein Structure Database^33^ and superimposed on HLS using Fr-TM-Align^34^. The average predicted Local Distance Difference Test (pLDDT) scores were 96 for committed OSCs and 95.2 for multifunctional OSCs (scores range from 1-100) (Supplementary Fig. 2a), indicating high confidence in the predictions. We also calculated B-factors for the OSC structures extracted from the AlphaFold Protein Structure Database using an all atom contact model^35^. The results provide additional support for increased flexibility in multifunctional enzymes (Supplementary Fig. 2b). The binding pocket residues were then extracted from the superimposed structures and the resulting sequences (Data S2) used to generate the network and tree shown in Fig. 2b. Visual inspection of the sequence similarity networks and phylogenetic trees suggested that OSC function was satisfactorily captured by the binding pocket residues (Fig. 2a,b). To test how well the binding pocket sequence captured functional information compared to the full sequence, parsimony scores were calculated using the primary enzyme function on the generated phylogenies (see Materials and Methods). OSCs that primarily synthesized a unique product (not observed in any of the other 169 OSCs) were labelled as ‘other’, and the difference in parsimony scores between the full-sequence and binding pocket trees when including or excluding these was not significant (excluding ‘other’ functions: *p* = 0.339, z =-0.591, including ‘other’ functions: *p* = 0.451, z =-0.258). Therefore, the binding pocket residues were at least as effective in capturing the sequence-function phylogenetic relationship as the full OSC sequences (Fig. 2c).

The 169 OSCs were all compared pairwise, with each comparison forming one of three sets for analysis according to their known product relationships: (i) comparisons between those that make different products; (ii) comparisons between those where at least one common product is shared, and (iii) comparisons between those that make only the same product (products listed in Data S1). Using this classification the average overall amino acid sequence identities for the full length sequences were 58%, 68% and 76% respectively, while for the binding pocket residues these identities increased to 63%, 81%, 94% (Supplementary Fig. 3a, b). We also compared the distributions of average Grantham scores of the aligned binding pocket residues. The Grantham score^36^ quantifies the ‘distance’ between amino acids based on their chemical properties (lower scores indicate more conservative substitutions). The mean Grantham scores for these OSC subgroups were 25, 13, and 4 respectively (Supplementary Fig. 3c). Together, these results show that enzymes already known to produce the same product exhibit higher sequence identity and more conservative bindingIZpocket substitutions, consistent with conservation of catalytic mechanisms.

### Investigation of the binding pocket residues of p-type OSCs

We next carried out an in-depth investigation of the likely product-determining residues of OSCs, focusing in on the three major groupings that make the protosteryl (p-type) products - cycloartenol, lanosterol and cucurbitadienol (Fig. 1b). These were selected because they form relatively closely related but distinct phylogenetic groupings (Fig. 2) and make different products, so presenting a tractable opportunity for deeper investigations of the amino acid residues that determine product specificity. These products represent alternative tetracyclic resolutions of the protosteryl cation, interrelated through a sequence of 1,2-hydride and alkyl shifts that progressively migrate the positive charge along the tertiary carbon backbone, with cucurbitadienol representing the most extensively migrated species (i.e. the largest number of shifts). The key cationic intermediates are the C9, C8, and C5 carbocations, which we will refer to as P9C, P8C, and P5C, respectively (Fig. 1b). A total of 36 cycloartenol synthase OSCs, three lanosterol synthase OSCs and five cucurbitadienol synthase OSCs were selected for this analysis (Data S1). For these OSCs, we compared AAindexIZderived descriptors (volume, hydrophobicity, flexibility)^37^ across the 50 binding pocket residues defined above (Supplementary Fig.□4). Although the three OSC subfamilies were represented by different numbers of sequences, the comparisons nonetheless revealed consistent trends in binding-pocket properties. The observed differences were primarily in the flexibility and hydrophobicity surrounding the A-C rings, suggesting subfamilyIZspecific tuning of bindingIZpocket chemistry around the product rings (Supplementary Fig.□4a). To further investigate the features of the binding pockets, we analysed their electrostatic properties in the three different product subfamilies using Adaptive Poisson-Boltzmann Solver (APBS)^38^. The binding pocket is in the interior of the protein in all OSC enzymes^28,30^. Whilst many residues lining the pocket are hydrophobic, the overall potential of the pocket is predominantly electronegative. The pockets of both lanosterol-and cucurbitadienol-synthesising OSCs were found to be highly electronegative, while in those that make cycloartenol the pocket was only electronegative/hydrophobic around ring A (Supplementary Figure 4b).

### Identification and functional analysis of product determining residues required for conversion of a cycloartenol synthase into a cucurbitadienol synthase

We next compared the conserved residues identified in the 36 cycloartenol-producing enzymes with those identified in five cucurbitadienol enzymes (Data S1) by multiple sequence alignment using MAFFT^39^, and used the GroupSim program^24^ within the FunFamer protocol^23^ to identify differentially conserved residues between the two subfamilies. To evaluate confidence in the assignment of conserved residues, we considered the sequence diversity of the enzymes in each subfamily. This is reflected in the Diversity of Position (DOP) scores^40^. The cycloartenol-and cucurbitadienol-producing enzyme sets comprise highly diverse sequences with DOP scores of 79 and 84 respectively (out of a maximum of 100).

We identified four differentially conserved residues that distinguished the cycloartenol-and cucurbitadienol-producing enzymes (Y118L, I365L, T531S, S609C), together with two additional residues (G116S, P480L) located within 5 Å of these (secondary shell residues) (Fig. 3a) (AtCAS numbering is used throughout this study). The Groupsim score^24^, which quantifies differential conservation between groups, was ≥0.9 for all six positions, indicating strong group-specific conservation and evolutionary constraint. Grantham score analysis further showed that S609C represents a moderately radical mutation (>100 out of 215), whereas Y118L and I365L are conservative substitutions (< 50); the remaining substitutions were moderately conservative (50 - 100) (Fig. 3a). The differentially conserved residues form spatially clustered networks in the predicted 3D space, with each residue located within 5 Å of at least one other residue in the cluster. Residues 116 and 118 fall into one cluster, and the remaining four residues into another. The predicted product determining residues are in contact with rings A (480, 531 and 609), B (116, 118) and C (365) (Fig. 3b).

**Fig. 3.**
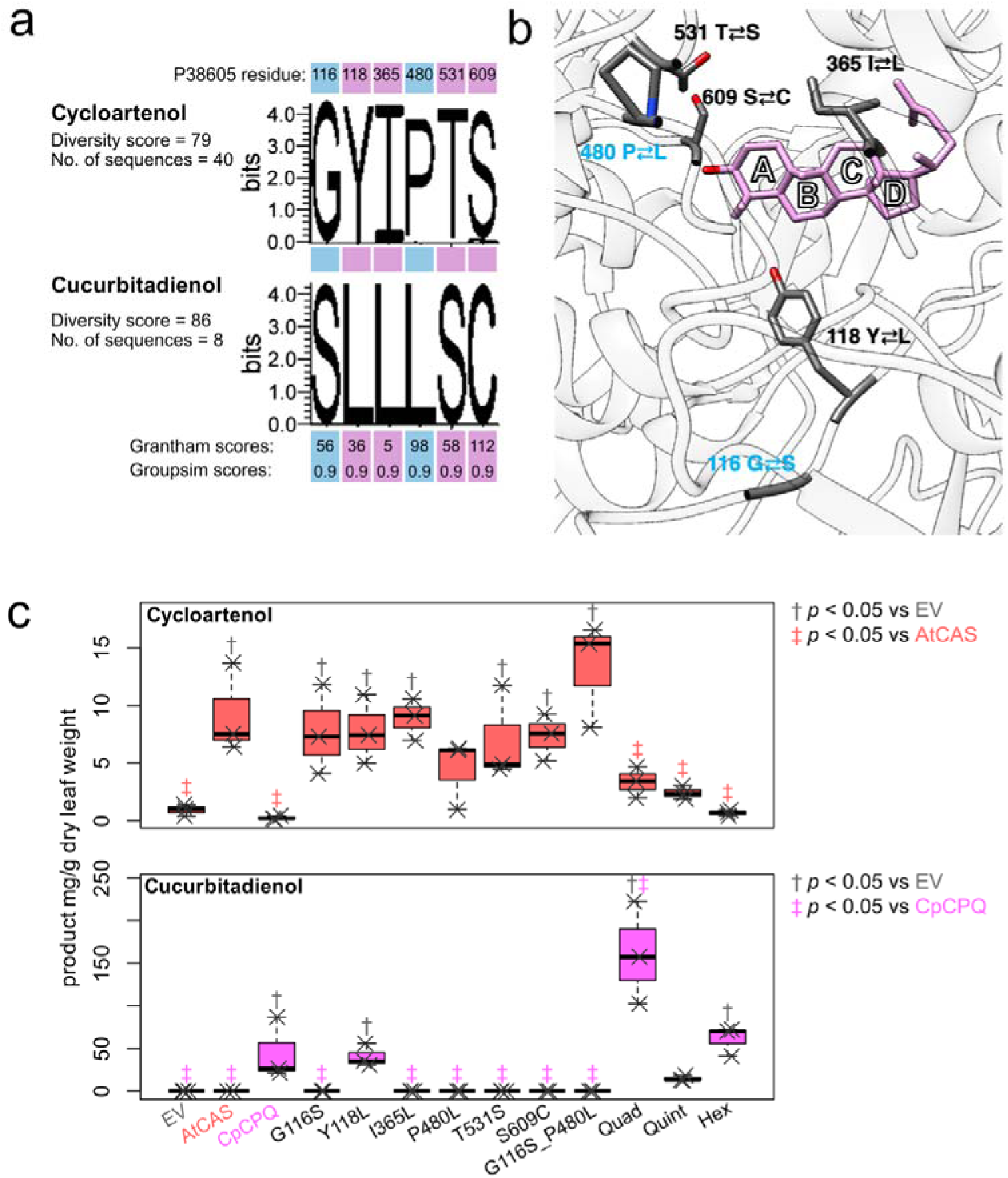
Identification and functional analysis of amino acid residues implicated in product specificity for either cycloartenol or cucurbitadienol. **a**, Differentially conserved binding pocket and secondary shell residues (highlighted in pink and blue, respectively) when cycloartenol-and cucurbitadienol-producing OSCs are compared, showing the DOPS diversity score and the Grantham and Groupsim scores for aligned residues. Residue numbering (based on the *Arabidopsis thaliana* cycloartenol synthase AtCAS; UniProt no. P38605) is shown at the top. **b**, Location of the differentially conserved binding and secondary shell residues. Residues 480, 531 and 609 are within 5 Å of each other, and residues 116 and 118 are also within 5 Å of each other. **c**, GC-MS quantification of cycloartenol and cucurbitadienol levels (a minimum of 3 biological replicates/treatment). Quantification was carried out as described in the Methods section. Post-hoc Dunnett’s test was carried out to assess the change in yields vs empty vector (EV) control (†) and wild type (AtCAS/CpCPQ) controls (‡) (see Data S3).

Following identification of the amino acid target residues implicated in determining product specificity for either cycloartenol or cucurbitadienol, we then carried out mutational analysis to evaluate the roles of these residues and to establish whether we could convert a cycloartenol synthase to a cucurbitadienol synthase. To do this we selected *Arabidopsis thaliana* cycloartenol synthase (AtCAS)^41^ as our template and generated a series of single and multiple mutant variants for the six amino acids identified above (Table 1). These mutant variants were cloned into the pEAQ-HT-DEST1 vector^42^ for functional evaluation by *Agrobacterium*-mediated transient expression in *Nicotiana benthamiana*, a well-established heterologous expression system for functional analysis of plant OSCs^10,43^. The wild type *A. thaliana* cycloartenol synthase AtCAS^41^ and a previously characterized cucurbitadienol-producing OSC from *Cucurbita pepo* (CpCPQ; ∼68% amino acid similarity cf. AtCAS, Supplementary Table 1)^44^ were included as positive controls. Five days after agro-infiltration, leaves were harvested and their extracts analysed by GC-MS (Fig. 3c; Supplementary Figures 5 and 6). The majority of the single mutants remained ‘committed’ cycloartenol synthases. However, the Y118L mutant was able to produce cucurbitadienol at a similar level to the wild type CpCPQ enzyme, while still retaining its cycloartenol synthase activity, making it a multifunctional OSC (Fig. 3c). The double mutant G116S_P480L (both secondary shell residues) did not produce cucurbitadienol, thus these two residues do not determine specificity for cucurbitadienol as the product. Interestingly the multiple mutants that also contain the Y118L mutation show different and contrasting product profiles. The quadruple mutant AtCAS_Quad (Y118L_I365L_T531S_S609C) produced substantially more cucurbitadienol than the Y118L mutant, while still producing some cycloartenol. Thus one or more of I365L, T531S, and S609C must act in combination with Y118L to favour enhanced cucurbitadienol production. The quintuple mutant AtCAS_Quint (G116S_Y118L_T531S_P480L_S609C) was substantially compromised in cycloartenol production and made little/no cucurbitadienol; while the hextuplet mutant AtCAS_Hex (G116S_Y118L_I365L_T531S_P480L_S609C) produced cucurbitadienol at a similar level to the wild type CpCPQ enzyme and had lost the ability to produce cycloartenol, i.e. has a switched product profile (Fig. 3c). We have already shown through analysis of the double mutant G116S_P480L that the two secondary shell residues do not determine product specificity for cucurbitadienol. Collectively, these results suggest that Y118L is a permissive change that enables cucurbitadienol formation, and that I365L enhances the specificity of this outcome in multiIZmutant contexts (e.g., AtCAS_Quad, AtCAS_Hex). Our data support Y118L as necessary but not sufficient for WT cucurbitadienol activity. MD simulations suggest that the AtCAS_Y118L mutant has increased water accessibility to the C6 atom, similar to the wild type CpCPQ enzyme (Supplementary Figure 7); also that the AtCAS_Quad mutation, located in the binding-site pocket, produces greater structural perturbation in the immediate vicinity of the P5C ligand (Supplementary Figure 8).

**Table 1.** Wild type and mutant OSCs tested in this study. Products of enzymes are noted according to the following: –, no activity; ++++, greater than wild type activity; +++, wild type activity; ++, neither significantly different from the empty vector control or the wild type enzyme; +, possible activity but the statistical power of the post-hoc Dunnett’s test could not confirm it.

| OSC | Name | Product(s) |  |  |
| --- | --- | --- | --- | --- |
|  |  | Cycloartenol | Cucurbitadienol | Lanosterol |
| *Arabidopsis thaliana* cycloartenol synthase | AtCAS WT | +++ | – | – |
| *Cucurbita pepo* cucurbitadienol synthase | CpCPQ WT | – | +++ | – |
| *Arabidopsis thaliana* lanosterol synthase | AtLSS WT | – | – | +++ |
| AtCAS: G116S | G116S | +++ | – |  |
| AtCAS: Y118L | Y118L | +++ | +++ |  |
| AtCAS: I365L | I365L | +++ | – |  |
| AtCAS: P480L | P480L | ++ | – |  |
| AtCAS: T531S | T531S | +++ | – |  |
| AtCAS: S609C | S609C | +++ | – |  |
| AtCAS: G116S_P480L | AtCAS_Double | +++ | – |  |
| AtCAS: Y118L_I365L_T531S_S609C | AtCAS_Quad | + | ++++ |  |
| AtCAS: G116S_Y118L_T531S_P480L_S609C | AtCAS_Quint | + | ++ |  |
| AtCAS: G116S_Y118L_I365L_P480L_T531S_S609C | AtCAS_Hex | – | +++ |  |
| AtCAS: A469G | A469G | +++ |  | – |
| AtCAS: I481V | I481V | +++ |  | + |
| AtCAS: T531S | T531S | +++ |  | – |
| AtCAS: I481V_T531S | I481V_T531S | ++ |  | + |
| AtCAS: A469G_T531S | A469G_T531S | +++ |  | – |
| AtCAS: A469G_I481V_T531S | AtCAS_Triple | + |  | + |
| AtCAS: H477N | H477N | – |  | +++ |
| AtCAS: H477N_I481V | H477N_I481V | – |  | +++ |
| AtCAS: H477N_A469G_I481V_T531S | H477N_A469G_I481V_T531S | – |  | +++ |
| AtLSS: H257N | AtLSS:H257N | – |  | +++ |
| AtLSS: Y532F | AtLSS:Y532F | – |  | +++ |
| AtLSS: H257N_Y532F | AtLSS:H257N_Y532F | + |  | – |

### Identification and functional analysis of product determining residues required for conversion of a cycloartenol synthase into a lanosterol synthase

We next compared the 36 cycloartenol synthases with the three plant lanosterol synthases (Data S1). The DOP scores for these two groups were 79 and 39, respectively (out of a maximum of 100). Thus, the cycloartenol-producing enzymes have a highly diverse set of sequences, while the lanosterol-producing ones do not, despite being from different plant lineages (*A. thaliana*, *Panax ginseng,* and *Euphorbia lathyris*, from the Brassicaceae, Araliaceae and Euphorbiaceae families, respectively). We identified two binding pocket residues, I481V and T531S, as significantly differentially conserved between the cycloartenol and lanosterol-producing enzymes (Groupsim scores of 1.0 and 0.9, respectively) (Fig. 4a). Both of these residues are in contact with ring A (Fig. 4b). In addition, two secondary shell residues in contact (<5Å) with residue 481 also had Groupsim scores of greater than 0.7 (A469G and H477N) (Fig. 4b). H477N is a known mutation that changes a cycloartenol-producing enzyme to a lanosterol-producing one^45^. There were no other binding pocket residues with a Groupsim score of 0.7 and above. We therefore identified these four residues as likely to impact on the product. Grantham score analysis showed I481V as a conservative substitution (< 50) and the other residue substitutions as moderately conservative (50-100). Since published QM/MM studies have suggested that the H232 and Y503 residues are important for the quenching of the P9C/P8C intermediates (Fig. 1b) in a lanosterol-producing enzyme to produce lanosterol^46^ (H232 and Y503, equivalent to AtCAS residues 257 and 532), we also included these in our analysis.

**Fig. 4.**
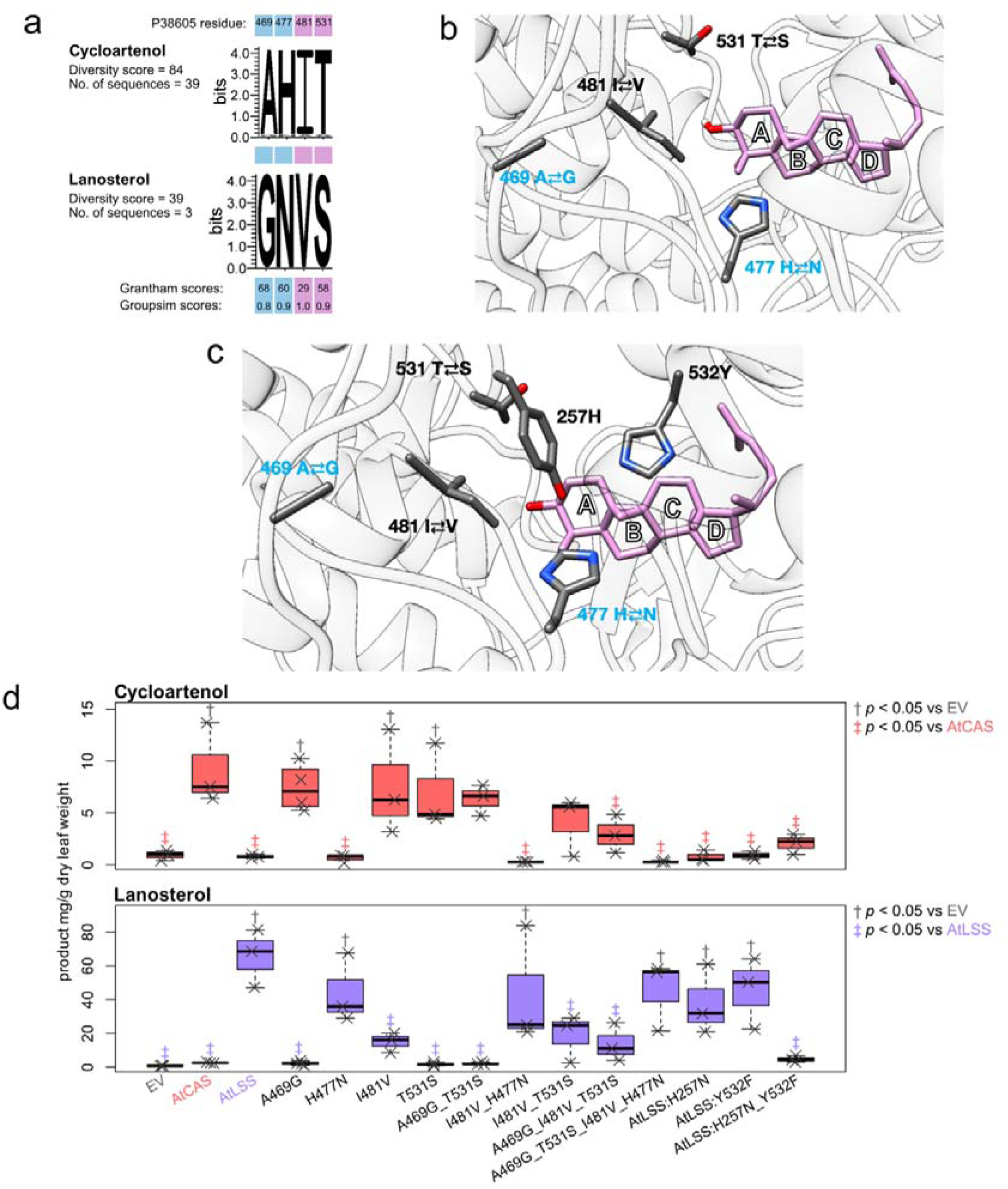
Identification and functional analysis of amino acid residues implicated in product specificity for either cycloartenol or lanosterol. **a**, Differentially conserved binding pocket and secondary shell residues (highlighted in pink and blue, respectively) when cycloartenol-and lanosterol-producing OSCs are compared, showing the DOPS diversity score and the Grantham and Groupsim scores for aligned residues. Residue numbering (based onAtCAS) is shown at the top. **b**, Location of the differentially conserved binding and secondary shell residues identified in this study. Residues 481, 531 and 469 are within 5 Å of each other. **c**, As for **b,** but also including residues 257 and 532 (residue numbering relative to AtCAS), which have previously been postulated (for AtLSS) to play a role in quenching the intermediate to produce lanosterol^46^. **d**, GC-MS quantification of cycloartenol and lanosterol levels (a minimum of 3 biological replicates/treatment). Quantification was carried out as described in the Methods section. Post-hoc Dunnett’s test was carried out to assess the change in yields vs empty vector (EV) control (†) and wild type (AtCAS/CpCPQ) controls (‡) (see Data S3). The two LSS mutants are shown on the right: H232 and Y503^46^ are equivalent to AtCAS residues 257 and 532.

A set of single and multiple AtCAS mutant variants was designed to test the roles of these residues in product specificity (Table 1). We also generated single and double mutant variants of *A. thaliana* lanosterol synthase (AtLSS; 65.1% sequence identity to AtCAS, Supplementary Table 1, Supplementary Figure 5) for the two residues implicated in quenching of the P9C/P8C intermediates (H257 and Y532), since these had not previously been tested experimentally (Table 1). These were evaluated by transient plant expression along with the wild type AtCAS and AtLSS OSCs. A single mutation, I481V, was sufficient for production of a significant level of lanosterol, although the mutant enzyme retained the ability to produce detectable levels of cycloartenol (Fig. 4d; Supplementary Figure 9). Mutation of AtCAS residues 469 and 531 did not have any significant effect, the mutant variants producing wild type levels of cycloartenol and no detectable lanosterol. The single mutation AtCAS_H477N, however, results in a complete switch in product specificity from a committed cycloartenol synthase to a committed lanosterol synthase, as previously reported^45^. Double and triple mutant AtCAS variants were also tested, the results overall confirming the partial and full effects of mutations I481V and H477N on product specificity, respectively. Mutation of either H257 or Y532 in AtLSS, which have previously been suggested to improve quenching of the P9C/P8C intermediates important for production of lanosterol (Fig. 1b)^46^ did not abolish lanosterol synthase activity. However the double mutant H257N_Y532F failed to produce lanosterol (Fig. 4d, Data S3).

## Discussion

Here we deploy computational biology approaches to predict determinants of product specificity in the OSC enzyme superfamily. We then focus in on identification and experimental validation of product-determining amino acid residues for the ‘p-type’ OSCs, specifically those that make the protosteryl-derived triterpenes cycloartenol, cucurbitadienol or lanosterol. Using a multifaceted approach combining differential conservation along with structural information, physico-chemical properties of amino acids and electrostatics of the binding pocket we were able to understand differences in the binding pocket characteristics of the protosteryl product-producing enzymes. The differential conservation approach using GroupSim provides a statistically robust way of identifying residues which might be important for the product specificity of each enzyme type. Previous mutational studies of OSCs while extensive have been carried out on an *ad hoc* basis to investigate the properties of a given enzyme of interest. We therefore used the *A. thaliana* OSC AtCAS^41^ as a template for systematic mutagenesis with the aim of converting product specificity to either cucurbitadienol or lanosterol, so providing more comprehensive and directly comparable insights into the amino acid residues that determine product outcomes.

The predicted variants were tested by *Agrobacterium*-mediated transient expression in *N. benthamiana,* followed by mass spectroscopic quantification of the products produced. We were successful in converting the wild-type *A. thaliana* OSC AtCAS to produce either cucurbitadienol or lanosterol. Mutant Y118L produced cucurbitadienol but retained the ability to make some cycloartenol. The ‘Quad’ mutation, which had mutations at a further three binding site residues in addition to Y118L was the most effective in producing cucurbitadienol, most likely due to the increased volume of the binding pocket. For lanosterol, a single mutation in a binding pocket residue (I481V) was sufficient for some product conversion, again with residual ability to make cycloartenol, while a single amino acid change in a secondary shell residue (H477N) gave full conversion to lanosterol as a product. A summary of all of the key residues identified and tested in this study is provided in Supplementary Figure 10.

Building on these results, we sought to speculate how potential enzymatic reaction mechanisms might account for the differences in product profiles caused by the mutations studied, as well as the other physicochemical characteristics observed. Our hypotheses are summarised in Fig. 5 and described here. Firstly, it is well established that in animal LSSs the final β-elimination leading to lanosterol occurs with removal of a hydrogen from the C8 position of the C9 cation (P9C – in Fig. 5). To achieve this Tyr503 (AtCAS equivalent: Tyr532) facilitates the leaving hydrogen’s transfer to the basic His232 (AtCAS equivalent: His257) that acts as the final hydrogen acceptor^47^. Recent work^46^ has shown that in plant LSSs, a second tyrosine (Tyr410) can stabilize the preceding C8 cation (P8C) and sequester Tyr532, thereby disrupting the catalytic Tyr532-His257 dyad (AtCAS numbering) seen in animals. As a result, in plants, the final β-elimination instead sees a hydrogen removed from the C9 position of the C8 cation (P8C), catalyzed directly by His257, without the additional 1,2-hydride shift that occurs prior to the equivalent β-elimination from the C8 position (of the C9 cation - P9C) that is seen in animals (Fig. 5a).

**Fig. 5.**
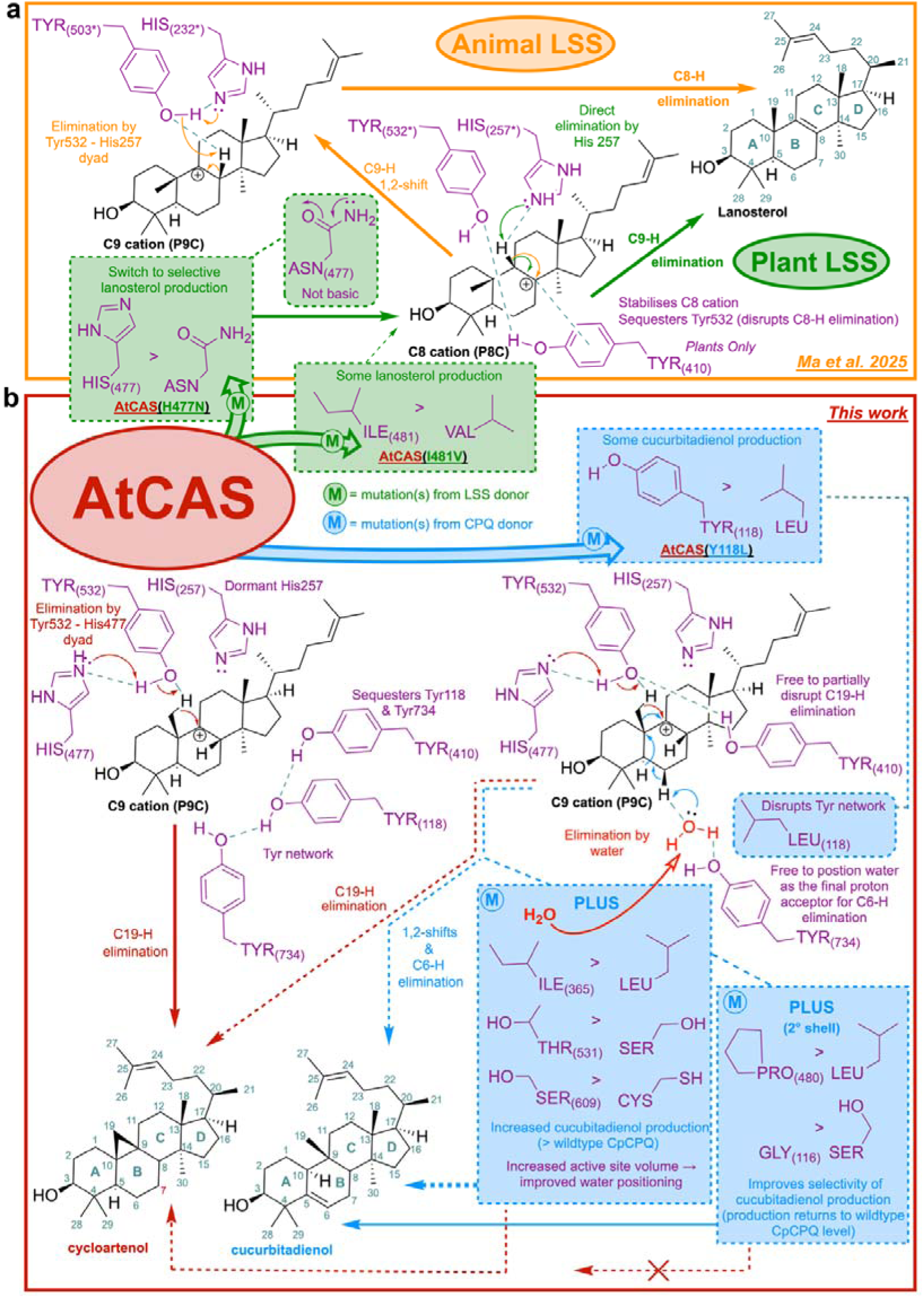
Proposed routes to lanosterol, cycloartenol and cucurbitadienol from the P8C and P9C cations, also summarizing the effects of the mutations studied in this work on the product profile of AtCAS. **a**, Comparison of animal and plant lanosterol synthases (LSSs), highlighting established differences in the catalytic site and mechanisms used for the final proton-elimination step in lanosterol biosynthesis based on Ma *et al* 2025^46^. N.B. Tyr503 and His232 in animal LSSs are equivalent to Tyr532 and His257 in plant LSSs. **b**, Proposed mechanism for cycloartenol formation by the plant cycloartenol synthase AtCAS and summary model illustrating how the key mutations characterized in this study impact on its product profile. Green shading = mutation from an LSS donor. Blue shading = mutation from a CPQ donor. Encircled M = mutation(s) made.

In plants, this C9 cation (P9C) is instead utilised in the formation of cycloartenol that requires elimination of a hydrogen from the vicinal C19 methyl group to yield the characteristic cyclopropane ring (Fig. 5b). In contrast, to yield cucurbitadienol this C19 methyl group itself shifts to the C9 cation as part of a cascade of additional 1,2-shifts that finally terminate in a β-elimination from the C6 position. Formation of the strained cyclopropane ring which terminates the cycloartenol pathway, would be expected to be significantly more energetically demanding, and less kinetically favorable than the two simple β-eliminations observed in the final steps of the lanosterol and cucurbitadienol pathways. This might be reflected by the unique asymmetric distribution of electrostatic potential we observed in the active site of cycloartenol synthases (Supplementary Figure 4); with this perhaps acting as a positive buffer to prevent retro shifting from the C9 cation (P9C) when formation of the cyclopropane is not kinetically favorable.

Our mutagenesis results align with previous studies showing that His477, found in plant cycloartenol synthases, is essential for cycloartenol formation^45^. This supports a model in which His477 partners with Tyr532 to eliminate a hydrogen from the C19 methyl group during formation of cycloartenol (Fig. 5b). When His477 is substituted with Asn (as in plant LSSs), the enzyme reverts to a committed lanosterol synthase. Although Asn477 cannot function as a final hydrogen acceptor (like His at this position), it can still likely donate a hydrogen bond and remain paired with Tyr532. This in turn allows the otherwise dormant His257 to resume its role as the direct base (seen in plant LSSs), restoring lanosterol formation via C9 hydrogen elimination (Fig 5a). Substitution of Ile481 with Val results in a small amount of lanosterol production. The side chain of residue 481 sits below His477, and this steric change may give His477 more freedom to align in a suboptimal position decreasing the efficiency of the His477-Tyr532 dyad (that is critical to cycloartenol formation) giving the opportunity for partial reversal to lanosterol synthase function (via direct His257 mediate hydrogen elimination).

Acquisition of cucurbitadienol synthase activity can be rationalised by evaluation of additional structural features within the active site of cycloartenol synthases that may further modulate these pathways (Fig 5b). In *A. thaliana* cycloartenol synthase (AtCAS), the Tyr410 (critical in plant LSS) is associated with two neighbouring tyrosines, Tyr118 and Tyr734, which together may form a stabilising hydrogen bond network. This network likely prevents Tyr410 from sequestering Tyr532, preserving the functional Tyr532–His477 dyad required for hydrogen elimination from the C19 methyl group and cycloartenol formation. When Tyr118 is replaced with Leu, this network is disrupted. Tyr410 becomes free to interfere with the dyad, and Tyr734 is no longer constrained. Under these conditions, Tyr734 may assist in positioning a strategically placed water molecule promoting a terminating β-elimination at C6 that results in the formation of cucurbitadienol, in addition to some residual cycloartenol (Fig. 5b). This is supported by MD simulations of wild type AtCAS, wild type CpCPQ, and AtCAS_Y118L that show improved occupancy of water near the C6 carbon for AtCAS_Y118L compared to wild AtCAS, making it more similar to wild CpCPQ (Supplementary Figure 7). The observation that additional replacement of active site residues Thr531, Ser609 and Ile365 with Ser, Cys, and Leu respectively in the AtCAS_Quad mutant, collectively (but not individually) dramatically increase cucurbitadienol yields, leads us to speculate that these changes may result in enhanced positioning to the C6 elimination site (especially considering the polar nature of two out of three residues involved in those mutations).

Selectivity in cucurbitadienol production is improved at the expense of increased yield, with collective addition of two substitutions, Pro at 480 for Leu, and Gly at 116 for Ser (secondary shell residues) (Fig. 3c). Even though the Pro to Leu and Gly to Ser substitutions lie outside the catalytic site, both could strongly influence the local fold and dynamics that govern substrate conformation, carbocation stability, and the timing of rearrangements. Proline strongly enforces backbone rigidity and is a classic “turn-promoting” residue, and glycine is a uniquely flexible one. Together such changes could reshape the substrate binding pocket remotely, and re-stabilize the backbone in a new geometry which completely disrupts the residual C19 hydrogen elimination, whilst the kinetically favorable C6 water mediated β-elimination can still occur (Fig 5b).

In summary, by combining a comprehensive comparative analysis with targeted bench validation of mutational sets in the same AtCAS background, we have moved beyond studyIZbyIZstudy examples of the OSC sequence-function relationship. We present an empirically supported framework for understanding and rewriting product specificity in plant OSCs. Together with the bindingIZpocket residue map, differentialIZconservation scoring, and physicochemical analyses, these results establish a practical blueprint for systematic, dataIZdriven exploration across this diverse, multiIZproduct enzyme superfamily. The unified strategy pinpoints and functionally confirms key product determining residues, and shows how cumulative, contextIZsensitive substitutions within a single reference sequence modulate both function and yield. Standardizing analyses and assays in the AtCAS background makes effects directly comparable. While tuning of dIZtype selectivity will likely demand broader remodelling, the same predictIZmutateIZmeasure workflow, executed efficiently in *N. benthamiana*, provides a clear path forward to build a coherent, testable program for predictive OSC engineering.

## Materials and Methods

### Collection and initial analysis of plant OSC sequences

A total of 169 OSC sequences were collated from the literature (Data S1), using the Trefor database^8^ to aid in selection. All-vs-all searches were carried out using MMSeqs easy-search^48^, and resultant identities were used as edge weights to plot a sequence similarity network using Gephi^49^ with the top 15% of edges displayed and ForceAtlas2 layout. A multiple sequence alignment and neighbor-joining (NJ) cladogram was generated using Clustal-Omega (https://www.ebi.ac.uk/jdispatcher/msa/clustalo) and simple phylogeny (https://www.ebi.ac.uk/jdispatcher/phylogeny/simple_phylogeny) with distance correction.

### Identification, extraction and initial analysis of binding pocket sequences

To define the binding pocket residues, we used the available crystal structure of human lanosterol synthase (HLS, PDB ID: 1W6K) and bacterial squalene hopene cyclase (SHC, PDB ID: 1UMP)^30–32^. We superimposed both the structures and considered all the residues within 5Å of 2-azasqualene in both HLS and SHC. Additionally, if any residues of a loop were within 5Å, the entire loop was considered as binding pocket residues because the loops could be flexible and the residue positions could be slightly different for other enzymes. This resulted in a total of 50 residues. 169 predicted structures for the functionally characterized plant OSCs were downloaded from the AlphaFold Protein Database^33^ and were superimposed on HLS using Fr-TM-Align^34^. The binding pocket sequence alignment was then extracted from the superimposed structures.

A sequence similarity network and a NJ cladogram was generated for the binding pocket residues as above. To test whether functional information was retained when only using the binding pocket residues, phylogeny-trait association was tested on the two tree topologies using Bayesian tip-association significance testing^50^. Briefly, this uses clustering of discrete tip states against null label permutations, whilst pairing permutations across trees to enable between-tree tests. Multifunctional OSCs were classed according to primary function, and OSCs producing a unique primary product in the set of 169 were categorized as ‘other’ (Data S1). Each labelled function was treated as a discrete category. For each tree, a parsimony score (PS) was calculated using the Fitch parsimony algorithm^51^, and one-sided permutation tests (with an add-one estimator) were used for significance, with 2000 permutations. Per-function *p-*values were adjusted using Benjamini-Hochberg false discovery rate correction^52^, and z-scores reported relative to null. This was carried out with the full set of OSC sequences (169 tips and 13 functions; full length vs. binding pocket ΔPS =-2, one-sided *p* = 0.339, z =-0.591) and without sequences classified as ‘other’ (150 tips and 12 functions; full length vs. binding pocket ΔPS =-1, one-sided *p* = 0.451, z =-0.256).

### Conservation of residues across the OSC enzyme family

We calculated the sequence identity between each aligned enzyme based on: (A) the whole protein sequence; and (B) just the binding pocket residues. We compared the sequence identity between: (i) all enzymes that produce different products; (ii) enzymes that produce at least one common product (i.e. committed and multifunctional enzymes); and (iii) committed enzymes that produce the same single product. Additionally, we calculated the similarity in physico-chemical properties of aligned amino acids residues in the binding pocket, using the Grantham score^36^. In order to study the conservation at each residue position in the OSC enzymes we first used MAFFT^39^ to create a multiple sequence alignment (MSA) of relatives in each subfamily of OSC enzymes. For each product type we created two different subfamilies. One subfamily containing just the committed enzymes, and another containing both committed and multifunctional enzymes. We then used Scorecons^40^ to identify the conservation score at each position of the MSA. We compared the distribution of conservation scores in the binding pocket residues, between committed and multifunctional enzymes.

### Identification of residues closer to each ring of the scaffold

Squalene in squalene hopene cyclase (SHC) was modelled using 2-azasqualene in the binding pocket (the folded substrate poised to make hopene), as reference. For this, the 2-azasqualene nitrogen is replaced with carbon and the bond is corrected according to the appropriate valency. A hopene-like conformation was generated by cleaving all ring bonds and reducing all double bonds to single bonds in Avogadro^53^.The resulting structure was subsequently subjected to geometry optimization to obtain a hopene-like fold. The hopene-like fold was superimposed on a squalene bound to SHC. Subsequently HLS was also superimposed on SHC to obtain a model of HLS bound to a hopene-like fold. The binding pocket residues were then sorted according to the proximity of the bound ligand in the hopene-like fold in Pymol^54^.

### Properties of the binding pocket residues

AAindex^37^ is a database that attributes numerical values to various physico-chemical properties of the amino acid. We used AAindex to calculate the properties of the binding pocket residues that are close to each of the rings. We calculated the hydrophobicity, flexibility and volume of the amino acid residues in the binding pockets of the OSC enzymes studied. In addition to studying the properties of the individual amino acids we also used Adaptive Poisson-Boltzmann solver (APBS)^38^ to calculate the electrostatics of the binding pocket for each of the product subfamilies. We used Chimera^55^ to create the surface representation of the proteins and then coloured the surface using electrostatic colouring based on the electrostatic potential calculated by APBS. We plotted the physico-chemical properties of the amino acids and the electrostatics of the binding pockets, for proteins in subfamilies producing lanosterol, cycloartenol and cucurbitadienol in order to identify differences in the binding pocket that might lead to different product formation. We used Tukey’s HSD to assess if the physico-chemical properties between the different product types were significantly different.

### Root Mean Square Deviation (RMSD) calculations of OSCs downloaded from the AlphaFold Database

The Cα atoms from each protein complex (optimised AlphaFold protein coordinates with intermediate P5C, see text) were superimposed using a least-squares fit, and the resulting pairwise RMSDs (Å) are shown in the symmetric matrix. These three closely related homologous plant proteins display low RMSDs, reflecting strong structural conservation. RMSD matrix derived from 6 Å binding-site environments following Cα-guided superposition. Protein structures were aligned using a Cα-only rigid-body fit, and all atoms within 6 Å of the P5C intermediate ligand were subsequently extracted to characterize the local binding-site environment. Pairwise RMSDs (Å) were computed to quantify pocket-level similarity.

### Predicted B-factor of the binding pocket residues

Predicted structures for the functionally characterized plant OSCs were downloaded from the AlphaFold Protein Database^33^, and used an all atom contact model to calculate the B-factors. The model combines features of the local density model (LDM) and the local contact model (LCM) for the prediction of B-factors of all atoms^35^. We also extracted the pLDDT values of the C-alpha residues from the downloaded AlphaFold structures. The crystallographic B-factor measures the displacement of atoms from their mean position, indicating their flexibility and dynamic properties. pLDDT (predicted local distance difference test) values of predicted AlphaFold models, which reflects the ‘local confidence’ of the model, were also used as a proxy for flexibility. The higher the pLDDT value, the lower the flexibility. There was not much difference in the median B-factor values between the multifunctional and committed enzymes. However, the overall distribution of B-factor values for the multifunctional enzymes is higher than for committed enzymes, suggesting that multifunctional enzymes are more flexible. Similarly, the distribution pLDDT values for binding pocket C-alpha residues of multifunctional enzymes is broader than for committed enzymes, supporting the increased flexibility of binding pocket residues in multifunctional enzymes. From the B-factor analysis and pLDDT scores, the binding pockets in the multifunctional OSCs tend to comprise more flexible residues than those with committed products.

### Identification of product determining residues

To identify differentially conserved product determining residues between subfamilies of OSC enzymes producing different products, (namely between cycloartenol and lanosterol subfamilies, and between cycloartenol and cucurbitadienol subfamilies) we created a multiple sequence alignment (MSA) for each of the product subfamilies using MAFFT as mentioned above. We considered only committed enzymes in our analysis. We used an entropy-based method, GroupSim^24^, to compare the alignments of the two subfamilies and identify differentially conserved residues (i.e. residues that are conserved at equivalent positions in the multiple sequence alignment but with different residues conserved in the two subfamilies being compared). For each column of the MSA, GroupSim measures the average similarity between pairs of residues for a given product subfamily, and the average similarities between product subfamilies. The average within-product subfamily type similarity subtracted by average between-product subfamily similarity is the column score that represents the likelihood of this position to be a PDR. Residues were predicted as PDRs if the Groupsim score was >=0.7. The diversity was measured by the DOP score which is returned from an analysis of the MSA using the program Scorecons^40^. The DOP score ranges between 0-100 and a threshold of 70 has been successfully used to indicate diverse alignments in other studies^25^. We only retained residues as PDRs if they were either in the binding pocket of the ligand or in contact with residues in the binding pocket of the ligand (secondary shell residues. We used Scorecons^40^ to identify residues that were conserved between product subfamilies.

### Molecular dynamics simulations

#### Preparation of starting structure complex

The AlphaFold model of cycloartenol synthase from plant *Arabidopsis thaliana* was taken as the starting structure for the simulation. The human lanosterol synthase (hLS) bound with lanosterol crystal structure (PDB ID: 1W6K) was taken as the reference structure for placing the intermediates P5C, P8C and P9C. The AlphaFold model structure was first superimposed onto the hLS structure. All the intermediates were then placed in the same position with similar orientations in the binding pocket as observed for lanosterol in hLS. This was achieved by using the RDkit library (www.rdkit.org) with the maximum common substructure (MCS) first determined between each intermediate and lanosterol.

Subsequently the intermediate structure was superimposed on lanosterol using the MCS. Mutant cycloartenol synthases were modelled using Pymol^54^ and intermediates placed in the binding pocket as described above for the wild type. Similarly, the AlphaFold model of cucurbitadienol synthase from *C. pepo* was taken as the starting structure for the simulation of the wild protein. Mutant cucurbitadienol synthase was modelled using Pinol. The intermediate P5C in cucurbitadienol synthase was placed in the binding pocket as described above.

#### Force-Field Parameters for the intermediates

The partial atomic charges of the intermediates P5C, P8C and P9C were calculated using the AM1-BCC charge model with the antechamber program of AmberTools^56^. All other parameters were assigned from the General Amber Force Field. AMBER topology/coordinate files were created using the AmberTools parmchk and tleap programs. These files were converted from AMBER to GROMACS format using the acpype python script^57^.

#### Molecular System Setup

The *pdb2gmx* module within the GROMACS package^58,59^ was used to generate topology and coordinate files for wild type cycloartenol synthase, mutant cycloartenol synthase, wild type cucurbitadienol synthase and mutant cucurbitadienol synthase using the AMBER99SB-ILDN force field^60^. The resulting topology and coordinates files were merged with those of the intermediates to obtain files for the corresponding complexes. The molecular systems were positioned at the center of a dodecahedral periodic box, solvated by adding water molecules, and neutralized by adding counterions with an excess of 0.150 M NaCl to mimic a physiological ion concentration. The TIP3P parameters were used for water molecules during the simulations.

#### MD Setup

MD simulations were performed using the GROMACS 2023.4 simulation package^58,59^. The energies of the molecular systems were minimized using the steepest descent algorithm to remove atomic clashes. The systems were heated from 0 to 300 K over 100 ps NVT simulations, followed by equilibration of densities at 1 atm pressure over 500 ps NPT simulations. In these simulations, all heavy atoms were restrained at the starting positions with a force constant of 1000 kJ mol^-1^ nm^-2^. These restraints were linearly removed during a subsequent 1 ns NPT simulation. The temperature and pressure were regulated using the Berendsen algorithm. Subsequently, 5 x *500 ns equilibrium simulations with random initial velocities were performed for the wild Cycloartenol synthase with P5C, P8C and P9C and for mutant cycloartenol synthase with P5C, P8C and P9C. Similarly, 5 x *500 ns equilibrium simulations with random initial velocities were performed for the wild type cucurbitadienol synthase and mutant cucurbitadienol synthases with P5C. The temperature and pressure were maintained at 300 K and 1 atm with 0.1 and 1 ps time constants using the v-rescale temperature and the Parrinello–Rahman pressure coupling method, respectively. The short-range nonbonded interactions were computed for atom pairs within a 14 Å distance. Long-range electrostatic interactions were calculated using the Particle-Mesh-Ewald summation method with fourth-order cubic interpolation and a 1.2 Å grid spacing. The time-steps during the simulations were 2 fs with all bonds constrained using the parallel LINCS algorithm.

#### MD Trajectory Analysis Minimum distance calculations

To calculate the distance between the water molecules and the C6 carbon from where elimination takes place to form the final product cucurbitadienol, trajectories were first concatenated from all five simulations at timesteps of 50ps. Then the pairwise distance was calculated using the GROMACS *gmx pairdist* command between water molecules and C6 carbon with hydrogens attached.

#### Clustering of trajectories

The *gmx_clusterByFeatures distmat* tool was first used to calculate a distance matrix. Then distances from this matrix were taken as features to cluster trajectories based on protein-ligand interactions. Only ring atoms and the groups attached to them in the intermediates were taken during both distance and clustering calculations. Clusters will have 1000 nearest structures to the central structure based on RMSD values for each cluster as output.

### Transient expression in *N. benthamiana*

Transient expression was carried out as described previously^10,43^. Wild type and mutant OSC sequences were synthesized (Twist Bioscience), cloned into the pEAQ-HT-DEST1 expression vector^42^ and transformed into *Agrobacterium tumefaciens* (strain LBA4404) for heterologous expression in *N. benthamiana*. *A.*□*tumefaciens* strains were streaked onto selective Luria–Bertani (LB) agar plates (50□μg□ml^−1^ rifampicin, 50□μg□ml^−1^ kanamycin and 100□μg□ml^−1^ streptomycin) and incubated overnight at 28□°C. These cultured streaks were used to individually inoculate 10□ml of selective LB medium (50□μg□ml^−1^ rifampicin, 50□μg□ml^−1^ kanamycin and 100□μg□ml^−1^ streptomycin) and incubated overnight at 28□°C with shaking. Cells were pelleted by centrifugation and resuspended in freshly prepared MMA buffer (10□mM 2-(*N*-morpholino)-ethanesulphonic acid pH 5.6 (KOH), 10□mM MgCl_2_, 150□μM 3′5′-dimethoxy-4′-acetophenone). The optical density of cells in MMA buffer at 600□nm was adjusted to 0.2 before the strains harbouring OSCs of interest were mixed in equal volumes with a strain harbouring a feedback-insensitive form of HMG-CoA reductase (tHMGR) for co-infiltration of plants with a needleless syringe^10,43^. Plants were harvested after 5 days in greenhouse conditions (25□°C with 16□h of lighting) post-infiltration. Harvested leaves were frozen and freeze-dried overnight before further use.

### Extraction and GC-MS analysis

Freeze-dried leaves were added to 1.5 ml centrifuge tubes with a tungsten carbide bead and ground at 1300 rpm for 1 min in a Geno/Grinder, producing a fine, dry powder. Dried *N. benthamiana* leaf powder (20 mg/sample) was weighed into 2 mL GC-MS autosampler vials (Chromex Scientific). Standard solutions of cycloartenol (Extrasynthese), lanosterol and cucurbitadienol (Osbourn lab) were made to a concentration of 1 mg/mL in ethanol and used to make eleven 2-fold dilutions, which when dried gave a calibration series from 100 μg down to ∼0.1 μg. These steps were performed using a Gerstel MultiPurpose Sampler (MPS) to minimize pipetting errors. A saponification solution (EtOH:H_2_O:KOH 9:1:1 v: v:w) was prepared containing the internal standard (coprostan-3□-ol, Sigma-Aldrich) to a final concentration of 10 μg/mL. Aliquots (500 μL) were pipetted into each sample vial (leaf powder and standards) using the Gerstel MPS. Samples were incubated at 65 °C for 2 h with intermittent agitation. 250 μL of H_2_O was added to each sample and vortexed before addition of 500 μL hexane. Samples were vortexed (10 s) and 10 μL of the upper hexane phase was transferred to a fresh vial and dried. Dried samples were derivatised using 50 µl 1-(trimethylsilyl)imidazole-pyridine reagent (Sigma-Aldrich) prior to GC-MS analysis.

GC was performed on an Agilent 7890B fitted with a Zebron ZB-5HT Inferno column (35 m x 250 μm x 0.1 μm, Phenomenex). Injections (1 μL) were performed in pulsed split mode (30 psi pulse pressure, 20:1 split ratio) or splitless mode depending on the concentration of the sample with the inlet temperature set to 250 °C. For the 15-minute method, the GC oven temperature gradient began at 170 °C (held 2 min) then ramped to 300 °C at a rate of 20 °C/min and held for 6.5 mins (for a total run time of 15 mins). For the 32-minute method, the temperature gradient began at 170 °C (held 2 min), then ramped to 290 °C at a rate of 6 °C/min and held for 4 mins, then ramped to 340 °C at a rate of 20 °C/min and held for 4 mins (for a total run time of 32 mins). The GC oven was coupled to an Agilent 5977 A Mass Selective Detector set to scan mode from 60 to 800 mass units (solvent delay 8 min) using electron impact (EI) ionisation running at 3.9 scans/sec. Data was analysed using the Agilent Masshunter Quantification software (Version 11). The quantifier ions used were 370.4 (coprostan-3□-ol), 408.4 (cycloartenol), 393.4 (lanosterol), and 408.4 (cucurbitadienol).

Quantification values were divided by the exact weight of leaf powder to calculate yield per g of dry leaf weight. To test whether quantified yields of the tested enzymes were significantly different of those from the empty vector or wild type controls, ANOVA and post-hoc Dunnett’s test was carried out. Raw data, *p*-values and confidence intervals for Figures 3 and 4 are provided in Data S3.

## Acknowledgements

We would like to acknowledge the following funding sources: Biotechnological and Biological Sciences Research Council (BBSRC) grants BB/V015176/1 (NS, RK, MS, JMT, ChrO, AO), BB/S020144/1 (ChrO, NS) and BB/S016023/1 (MS, AO), the Novozymes Prize 2023 (Novo Nordisk Foundation; AO), Wellcome Discovery Award #227375/Z/23/Z (RC, AO), the BBSRC Institute Strategic Programme Grant ‘Harnessing biosynthesis for sustainable food and health’ (BB/X01097X/1; ChrO, AO) and the John Innes Foundation (ChaO, AO). We thank JIC Horticultural Services for assistance with plant cultivation and the JIC Metabolomics and NMR platforms for assistance with instruments and method development, EMBL funding for JMT, including compute and storage

## Author contributions

RK performed bioinformatics, identified binding site residues, performed flexibility analysis, MD simulations and evaluted mechanistic possibilities. NS analysed conservation of residues, properties of binding pocket residues, identification of PDRs and MD simulations. JMT, ChrO and AO conceived and designed the project. NB performed substrate modelling. ChaO performed the bioinformatic and statistical analyses. RC carried out OSC cloning, generation of mutant variants, transient plant expression, metabolite analysis, and purification and NMR analysis of OSC products. MS advised on the mechanistic aspects of OSC function. All authors contributed to the preparation of the manuscript.

## Supplementary information

Supplementary Figures 1-10 and Supplementary Table 1

## Source data

Data S1: Table of the 169 functionally validated OSCs used in this study

Data S2: Extracted OSC binding pocket residues

Data S3: Quantification of enzyme products from agroinfiltration of *N. benthamiana*

Data S4: ANOVA and Tukeys HSD for Supplementary Figure 4

