## Supplementary Information for "Systematic prediction and functional analysis of amino acid residues determining product specificity in the plant oxidosqualene cyclase superfamily"

**Supplementary Figures and Tables**

| **a**  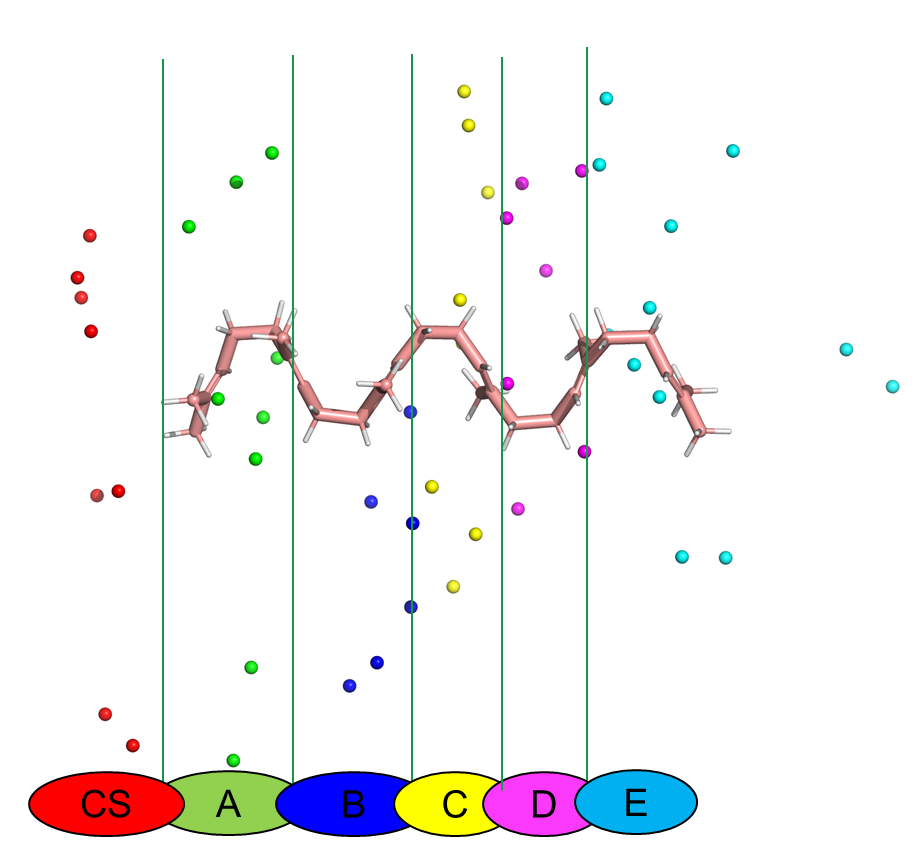 | **b**  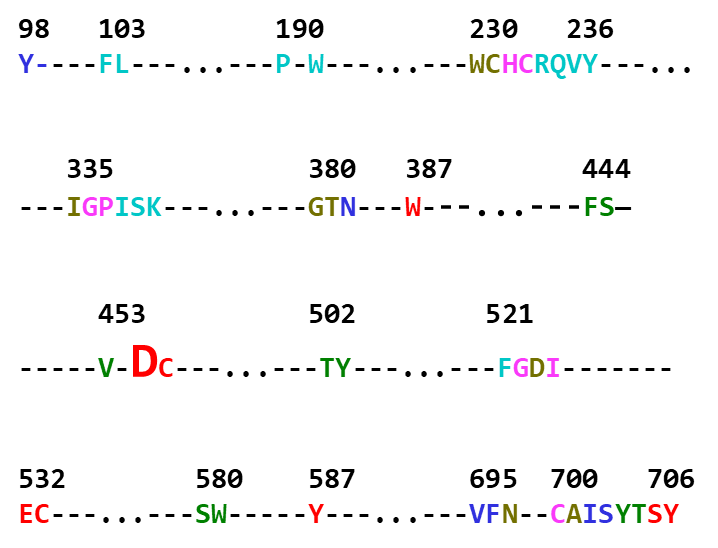 |
| --- | --- |

**c**

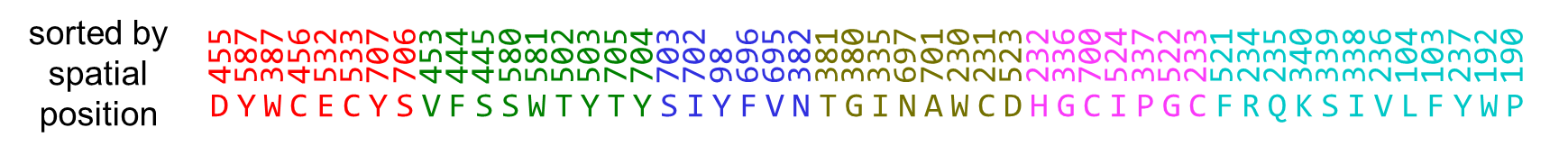

**d**

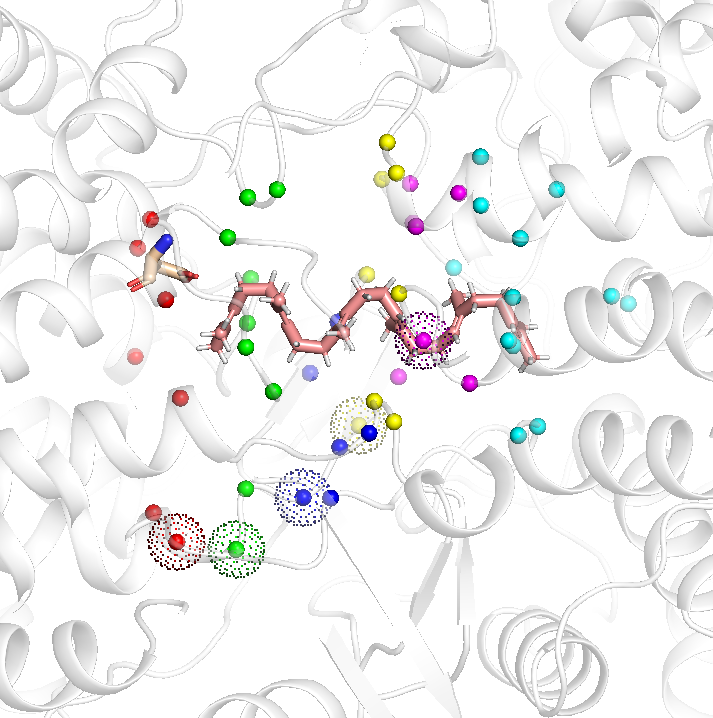

**Supplementary Fig. 1. Identification of binding pocket residues for HLS and SHC**

**a**, 3D spatial position of binding pocket residues for human lanosterol synthase (HLS) shown in 2D representation where the x-axis passes through the ligand and the y-axis denotes the coordinates of C-alpha atoms of binding pocket residues. Different colour balls represent the C-alpha atom of the catalytic site (CS) (red) - Catalytic site, A (light green) - 1^st^ ring, B (royal blue) - 2^nd^ ring, C (yellow) - 3^rd^ ring, D (magenta) - 4^th^ ring and E (cyan) - 5^th^ ring. **b**, Sequence position of binding pocket residues of HLS with catalytic residues shown in increased font. **c**, binding pocket residues arranged according to ring closeness. **d**, 3D spatial position of the binding pocket residues of HLS with respect to the ring-like configuration of squalene (there is no available crystal structure of OSCs with 2,3-oxidosqualene). Squalene is therefore modelled using squalene hopene cyclase (SHC) as a template. The SHC crystal structure is available with the inhibitor 2-azasqualene which is analogous to squalene and represents the open form of the substrate^28^. The catalytic residue ASP455 is shown in liquorice representation coloured by atom type. For all other residues the C-alpha atom is placed near the ring that it is closest to. CS (red) - Catalytic site, A (light green) - 1^st^ring, B (royal blue) - 2^nd^ ring, C (yellow) - 3^rd^ ring, D (magenta) - 4^th^ ring and E (cyan) - 5^th^ ring. If residues that are within 5 Å of squalene and lanosterol are also part of the flexible loop (700C, 701A, 703S, 705T and 706S), then the entire loop is considered as part of the binding pocket residues. The additional loop residues are highlighted using the dot representation.

| **a** 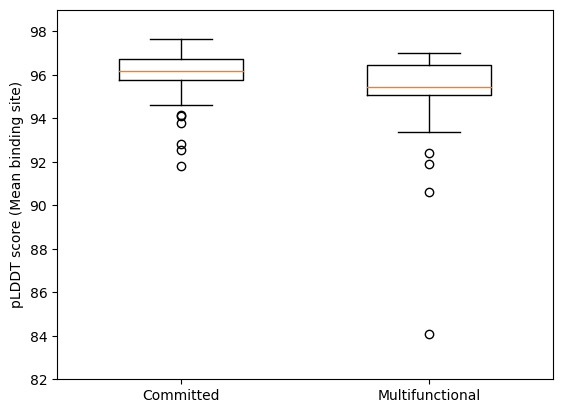 | **b**  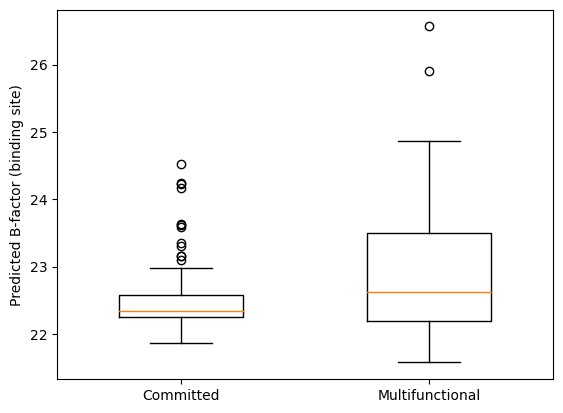 |
| --- | --- |

**Supplementary Fig. 2. Binding pocket residue analysis between multifunctional and committed OSCs**

**a**, Average predicted Local Distance Difference Test (pLDDT) plot for the binding pocket C-alpha atoms of 113 committed OSCs and 49 multifunctional OSCs (a total of 162 OSCs). The binding pocket has 50 residues. Boxes represent values between the 25th to 75th percentile and the orange line in the middle of the box represents the median value. **b,** Predicted B-factor plot for the binding pocket residues. To filter out the most flexible residues, the 90th percentile of all predicted b-factor values for binding pocket residues were calculated and are shown in the plot.

**a	Full length sequences	b	Binding pocket sequences**

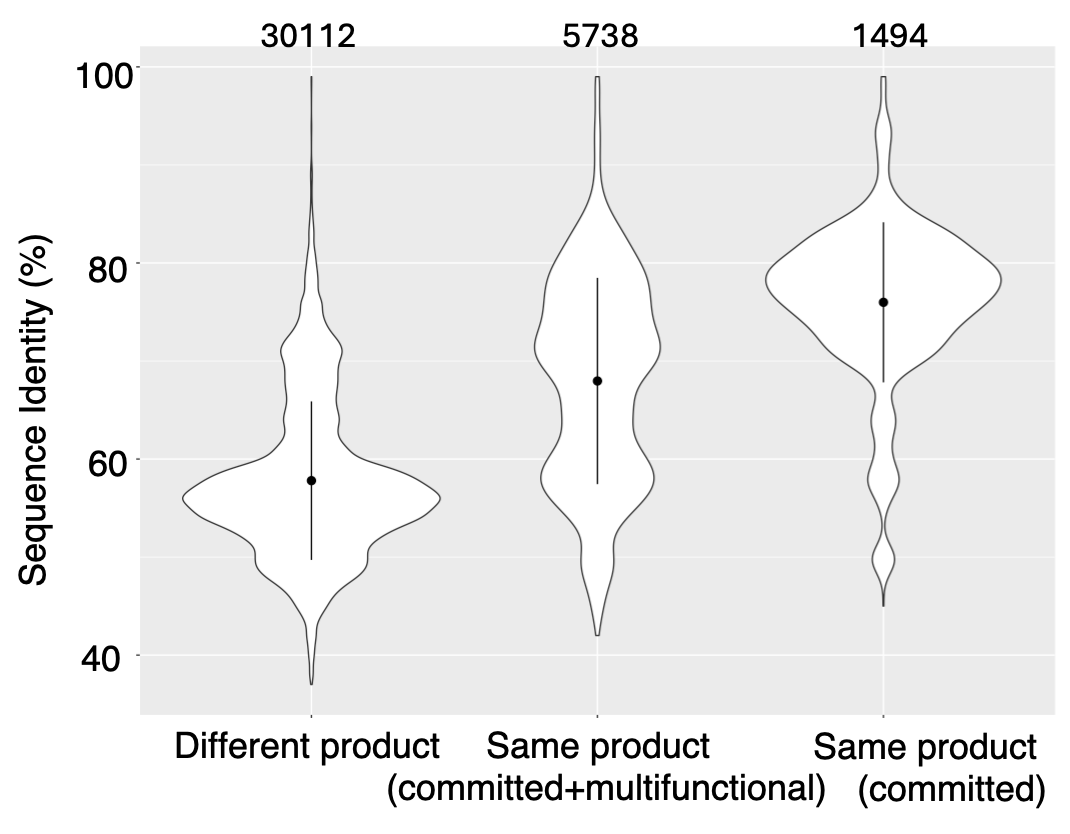

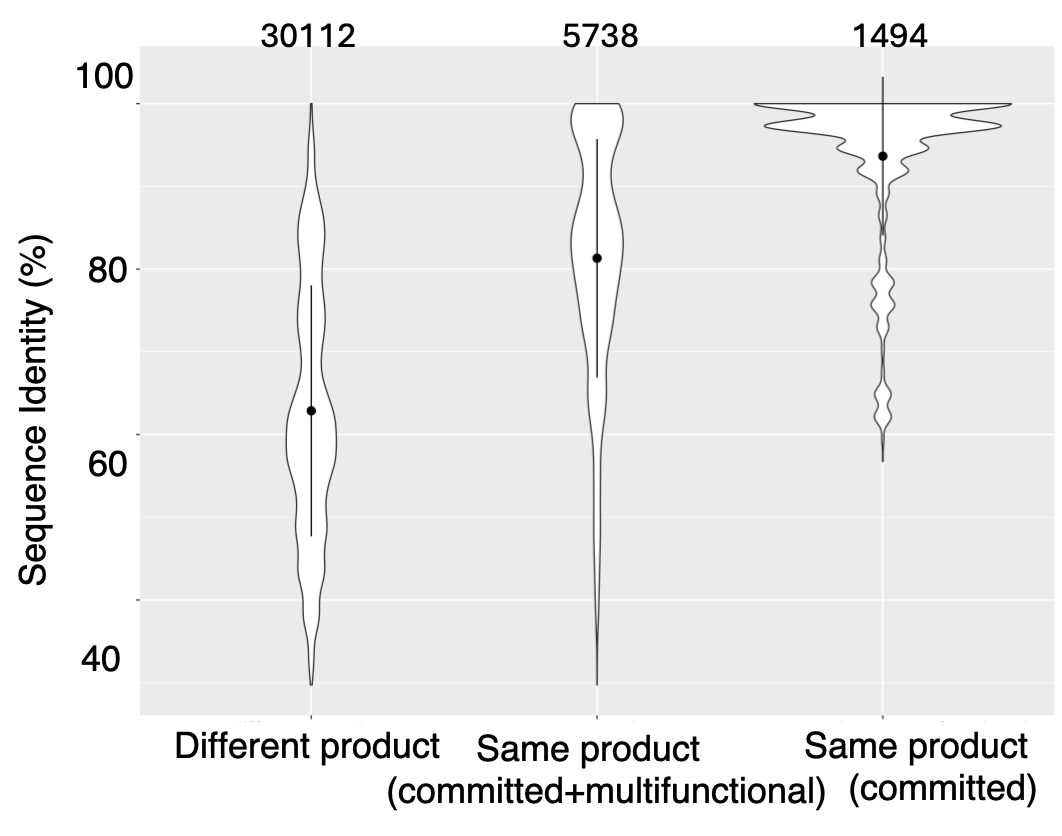

**c	Grantham scores**

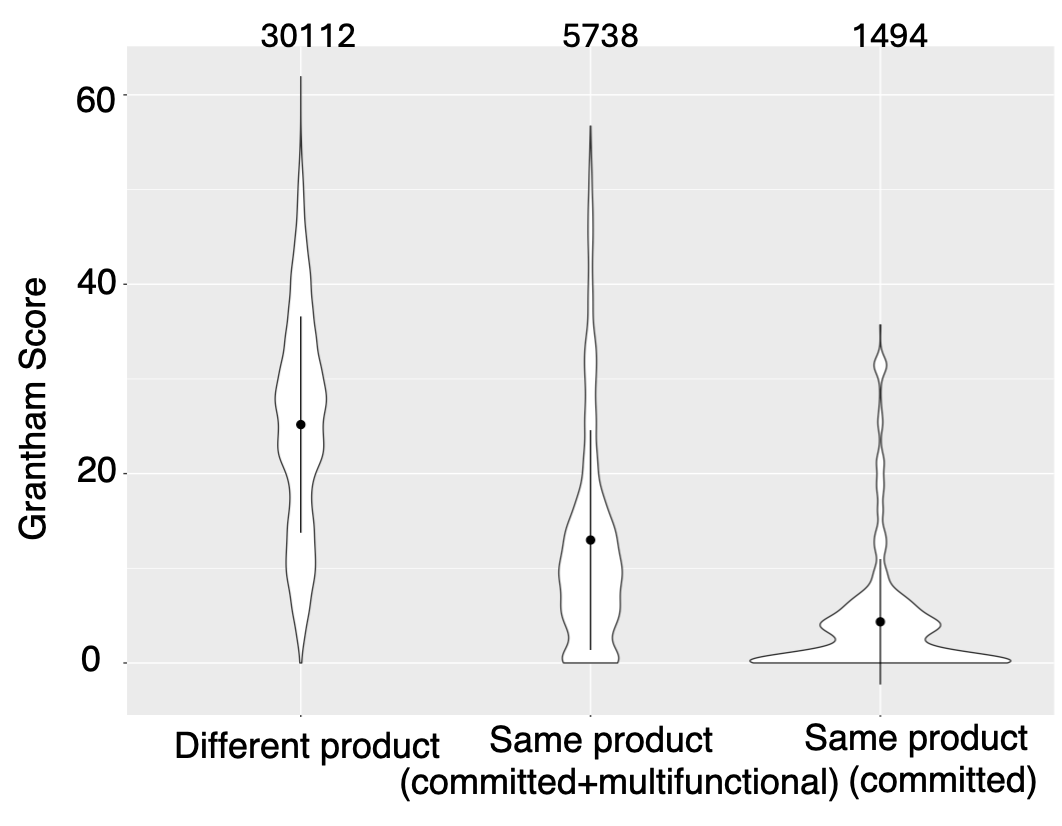

**Supplementary Figure 3. Analysis of amino acid sequence similarities for subgroups of OSCs that make different products, multiple products and the same product.**

**a**, Global protein sequence identities and **b**, binding pocket sequence identities between subgroups of OSCs that produce different products, the same product (committed and multifunctional) and the same product. **c**, Distribution of the average Grantham scores of the binding pocket residues for OSCs producing different products, same products (committed and multifunctional) and the same product. Mean values are indicated as a single point with standard deviation shown as a line. The total number of comparisons in each column of the plot are shown at the top of each panel.

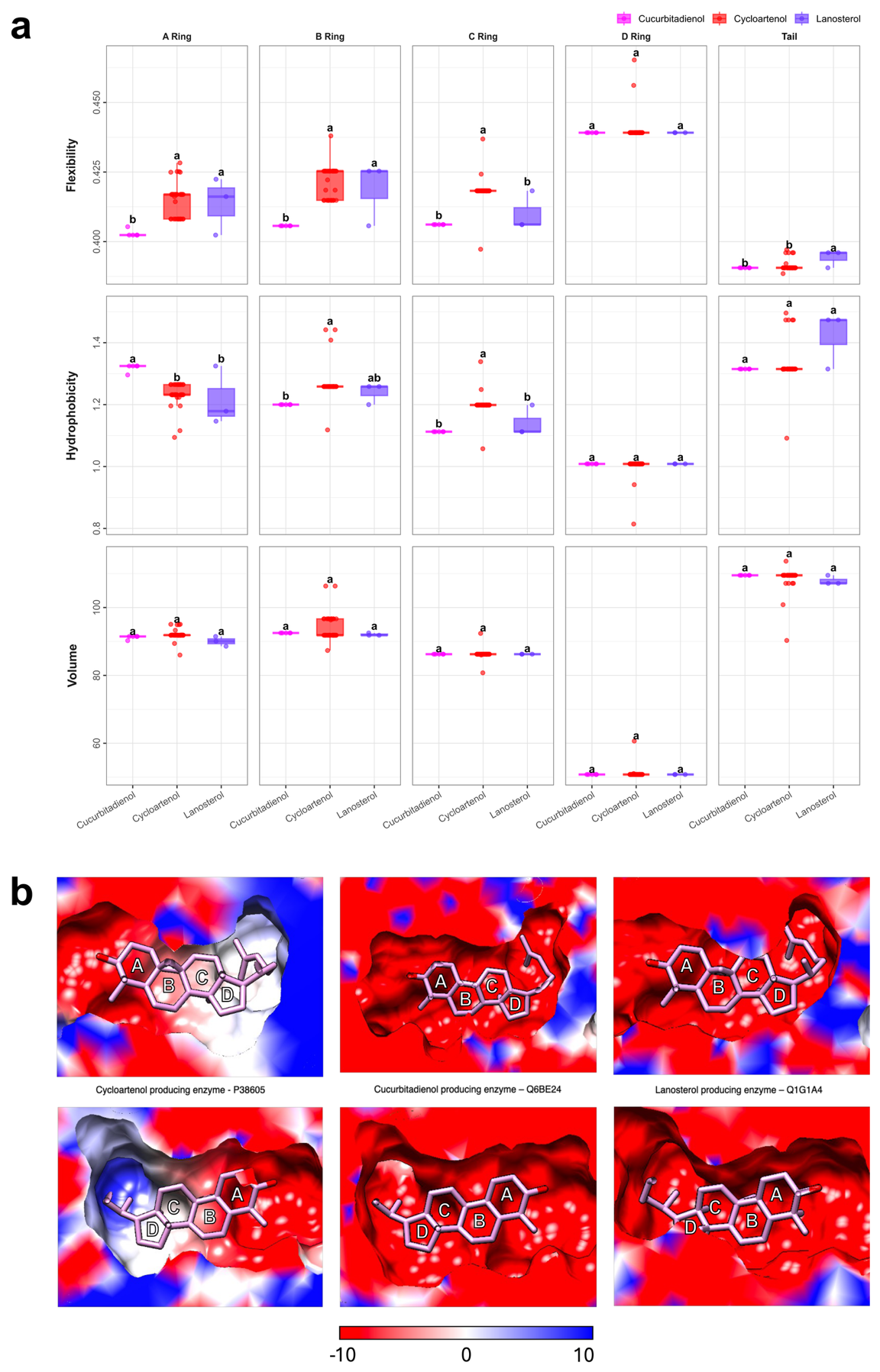

**Supplementary Figure 4. Analysis of the binding pocket residues for OSCs that biosynthesise protosteryl (p-type) products. a**, Flexibility, hydrophobicity and volume of the residues around each of the 4 rings and tail in cucurbitadienol-, cycloartenol- and lanosterol-producing OSCs (calculated using AAIndex and derived from the extracted binding pocket residues shown in Supplementary Figure 1). Different letters indicate statistically significant difference (p<0.05) within each section as calculated by Tukey’s HSD (Data S4) **b,** The binding pocket of cycloartenol-, cucurbitadienol- and lanosterol-producing OSCs coloured based on the electrostatics calculated using APBS: red, negative electrostatic potential; blue, positive electrostatic potential. The pocket is shown in two orientations 180 degrees to each other, rotated around the vertical axis in the plane of the ligand.

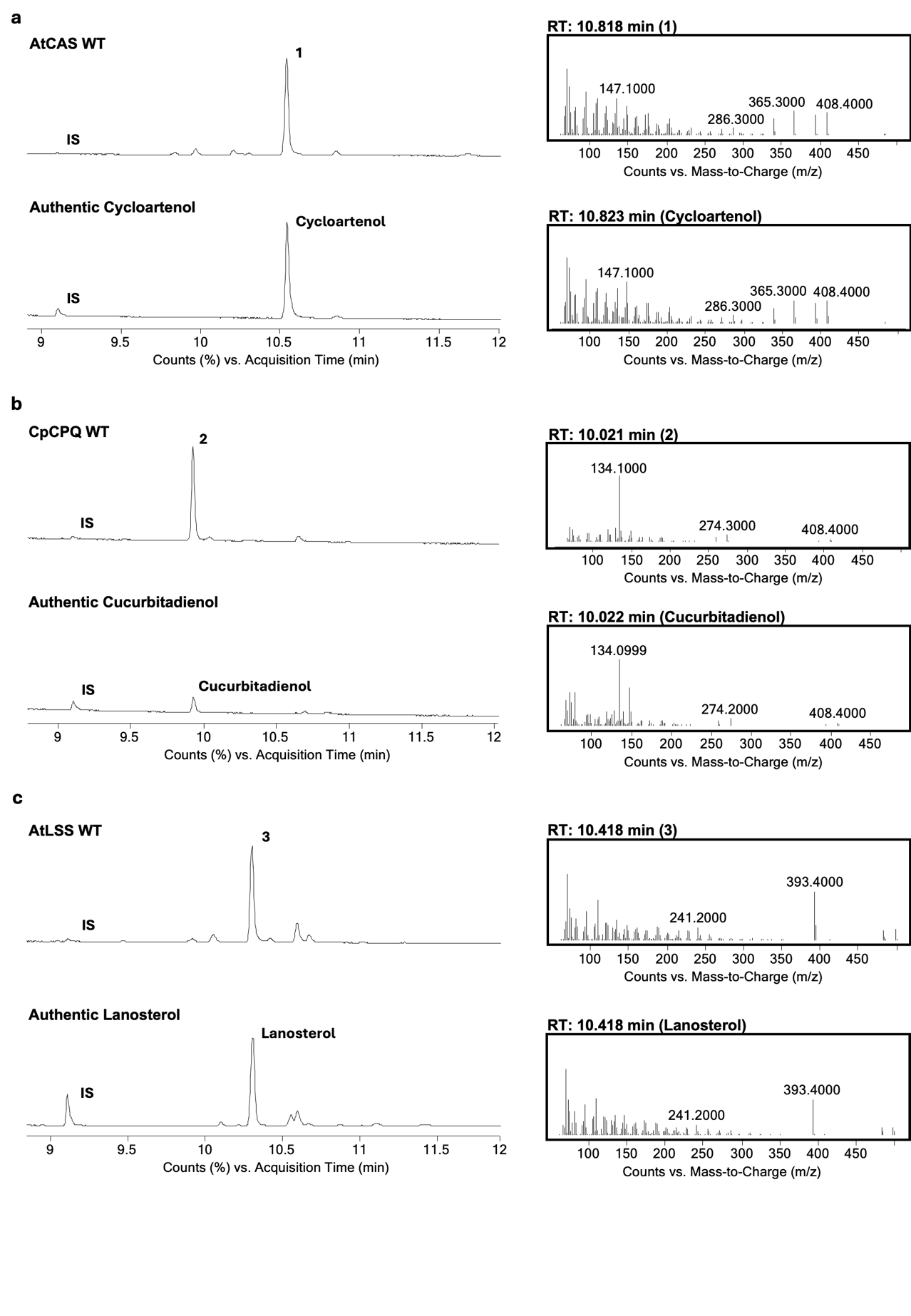

**Supplementary Figure** **5. Comparisons of wild type enzyme products and authentic standards.** Wild type enzymes were transiently expressed in *Nicotiana benthamiana* and leaf extracts analysed by GC-MS to assess the enzymatic product profile of: **a**, the AtCAS wild-type enzyme to an authentic cycloartenol standard; **b,** the CpCPQ wild type enzyme to an authentic cucurbitadienol standard; **c**, the AtLSS wild-type enzyme to an authentic lanosterol standard. Left: Total Ion Chromatogram, Right: Extracted Ion (EI) spectra. IS = internal standard, coprostanol.

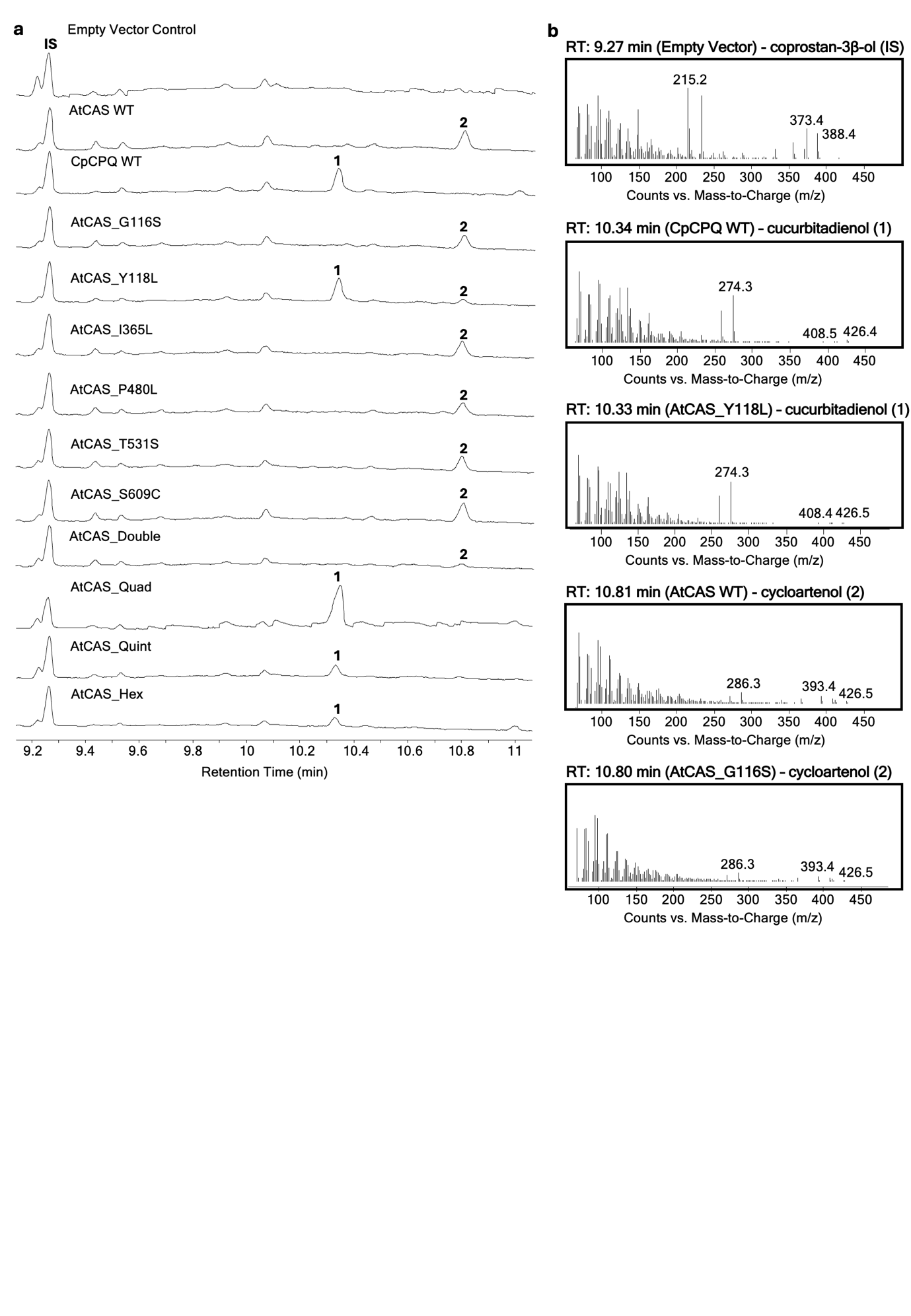

**Supplementary Figure 6. GC-MS analysis of *N. benthamiana* leaf extracts from plants transiently expressing wild type and mutant variants of AtCAS. a**, Representative chromatograms showing analysis of extracts from *N. benthamiana* leaves expressing wild type and mutant variants of *A. thaliana* cycloartenol synthase (AtCAS) or the cucurbitadienol-producing OSC CpCPQ. All enzymes were co-expressed with tHMGR. S = squalene, OS = oxidosqualene, IS = internal standard (coprostan-3β-ol, 20 µg/ml). **b,** +EI mass spectra of selected peaks.

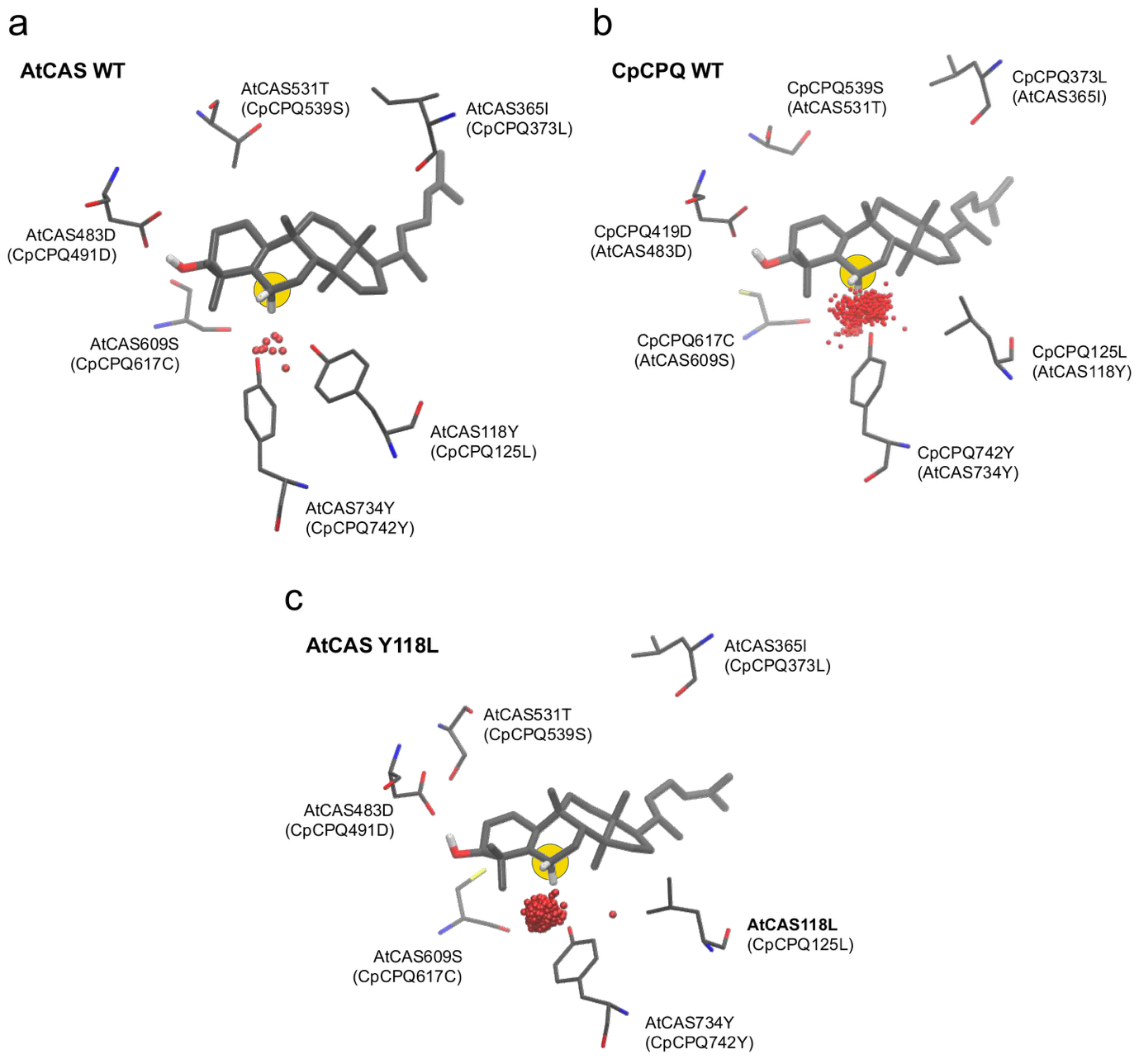

**Supplementary Figure 7. MD simulations of AtCAS and CpCPQ showing water occupancy at C6.** Occupancy of water molecules within 3 Å of the P5C C6 atom (yellow; see Figure 1 for numbering) is shown for the representative structure of the largest cluster for: **a**, AtCAS WT with P5C; **b**, CpCPQ WT with P5C; **c**, AtCAS Y118L with P5C. Water occupancy was calculated over 1000 snapshots from the cluster, whereas protein residues and ligand orientation are shown for a single representative snapshot for clarity. A clear difference is observed in the number of water molecules near the C6 carbon between wild-type AtCAS and wild-type CpCPQ. The AtCAS Y118L mutant shows increased water accessibility to the C6 carbon, similar to CpCPQ WT.

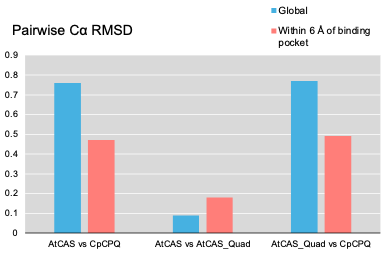

**Supplementary Figure 8. Pairwise Cα RMSD values derived from structural superposition and RMSD(Å) of all atoms (blue) and within 6 Å of the binding pocket of modelled P5C complex (red).** The four mutations introduced into wild-type cycloartenol synthase to produce AtCAS_Quad (Y118L, I365L, T531S, S609C) produce greater structural perturbation in the immediate vicinity of the ligand (red, centre) in comparison to the global Cα RMSDs (blue, centre). Conversely, the sequence-divergent wild type cucurbitadienol synthase (68% identity) exhibits a larger deviation at the global Cα level (blue, left) than within the local 6 Å binding-site region (red, left). AtCAS wild, *A. thaliana* cycloartenol synthase; AtCAS_Quad mutant (Y118L_I365L_T531S_S609C); CpCPQ, *C. pepo* cucurbitadienol synthase.

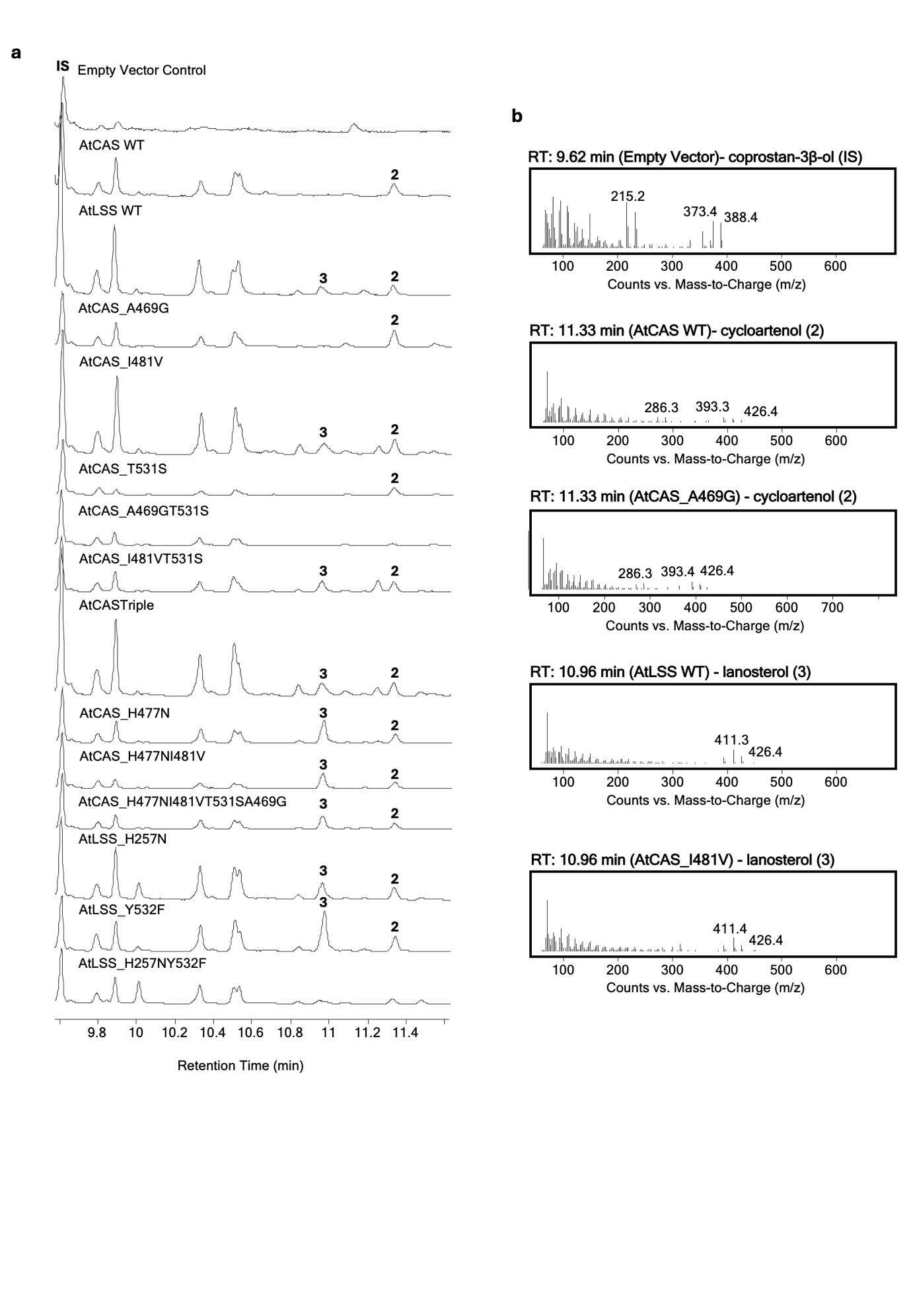

**Supplementary Fig. 9.** **GC-MS analysis of *N. benthamiana* leaf extracts from plants transiently expressing wild and mutant variants of AtCAS and AtLSS. a** Representative chromatograms showing analysis of extract from *N. benthamiana* leaves expressing wild type and mutant variants of AtCAS or AtLSS. All enzymes were co-expressed with tHMGR. IS = internal standard (coprostan-3β-ol 20 µg/ml) **b,** +EI mass spectra of selected peaks.

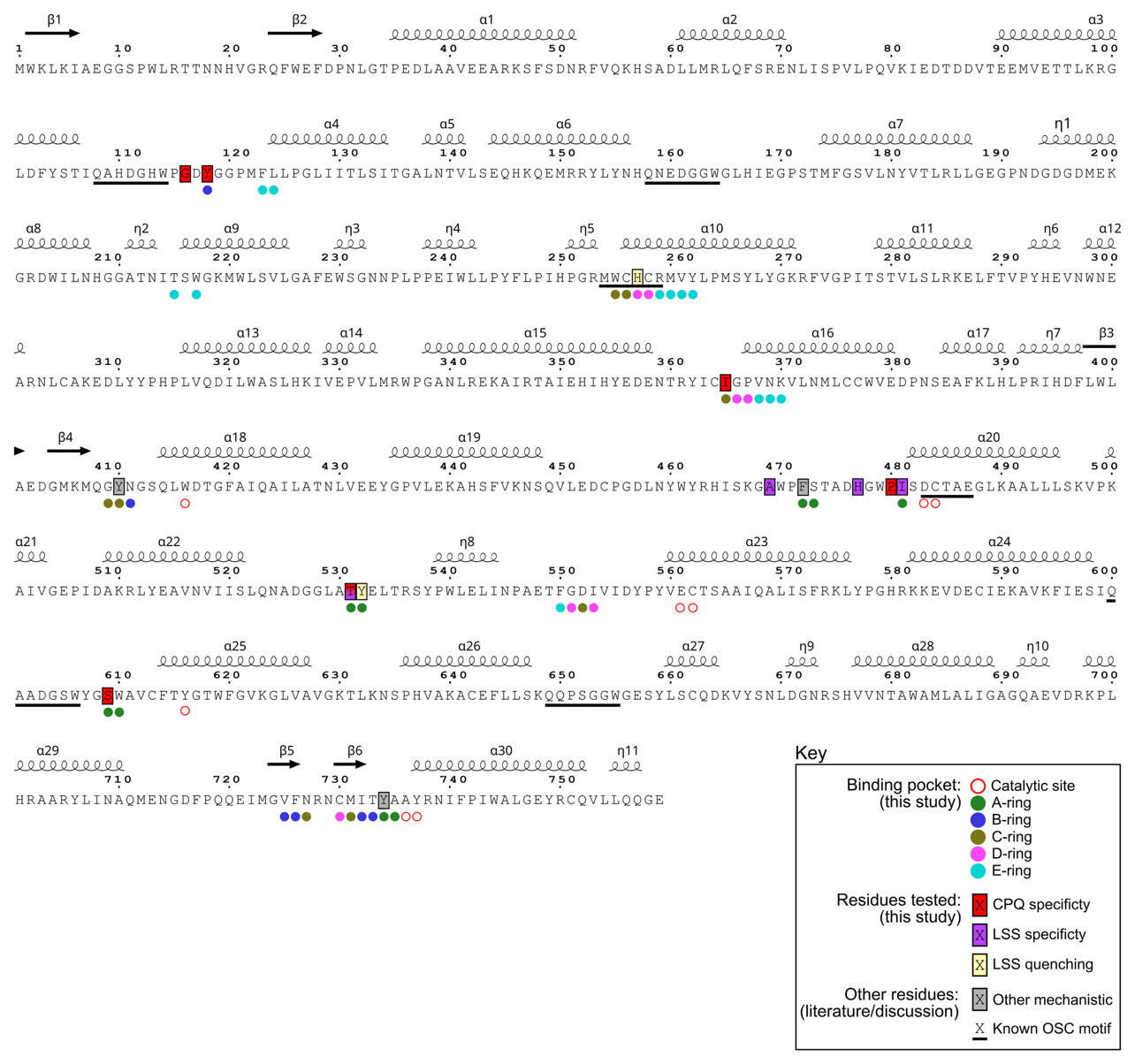

**Supplementary Fig. 10.** **Summary of key residues identified and tested in this study.** The AtCAS sequence is shown, with the secondary structural elements from the AlphaFold structural model overlaid above the sequence (generated with ESPript: <https://espript.ibcp.fr/ESPript/ESPript/>). Residues and motifs assessed in this study or discussed in the manuscript are labelled. Of the mechanistically relevant residues reported in the literature the only two not directly tested here are Y410^46^ (see discussion) and F472^61^, both of which are included as part of the proposed binding pocket residues. Also highlighted as ‘other’ is Y734, which we propose may have a role as part of a stabilizing hydrogen bond network (see Discussion).

**Supplementary Table 1. Pairwise amino-acid sequence identity matrix for five triterpene cyclases**. Protein sequences for cycloartenol synthase from *Arabidopsis thaliana* (AtCAS, P38605), cucurbitadienol synthase from *Cucurbita pepo* (CpCPQ, Q6BE24), lanosterol synthase from *Arabidopsis thaliana* (AtLSS, Q1G1A4), lanosterol synthase from *Homo sapiens* (LSS_HUMAN, P48449) and squalene–hopene cyclase from *Alicyclobacillus acidocaldarius* (SQHC_ALIAD, P33247) were compared using pairwise global alignment with the SIM alignment tool (ExPASy) https://web.expasy.org/sim. Percentage identity values shown represent the fraction of identical aligned residues relative to the alignment length excluding gap positions (Needleman–Wunsch scoring scheme, default parameters).

|  | **AtCAS** | **CpCPQ** | **AtLSS** | **LSS_HUMAN** | **SQHC_ALIAD** |
| --- | --- | --- | --- | --- | --- |
| **AtCAS** | - | 67.9 | 65.1 | 47.6 | 24.2 |
| **CpCPQ** | 67.9 | - | 65.5 | 47.2 | 33.3 |
| **AtLSS** | 65.1 | 65.5 | - | 46.8 | 24.7 |
| **LSS_HUMAN** | 47.6 | 47.2 | 46.8 | - | 25.3 |
| **SQHC_ALIAD** | 24.2 | 33.3 | 24.7 | 25.3 | - |
